# Evolutionary conserved reciprocal senescence and tumor suppressor signals limit lifetime cancer

**DOI:** 10.1101/2024.12.28.630560

**Authors:** Liqiong Liu, Stephani Davis, Shamila Yusuff, Ann Strange, Yonghua Zhuang, Prasanna Vaddi, Kathleen Flaherty, Kelly Jara, Joseph Kramer, Christine Archer, Schuyler Lee, Bifeng Gao, Adrie Van Bokhoven, Wei Wang, Sharon R. Pine, Tetsuya Nakamura, Hatim E. Sabaawy

## Abstract

Cellular senescence features a durable exit from the cell cycle triggered by stress or carcinogens. The INK4 locus is inactivated in various cancers, yet in senescence, p16^Ink4a^ is activated. Whether senescence is tumor-suppressing or -promoting remains a conundrum. We discovered an evolutionally-conserved Vertebrata INK4-homolog. This *ink4ab* triggers senescence upon oxidative- and/or carcinogenic-stress. Adult Ink4ab-deficient animals failed to activate senescence and developed spontaneous cancers. Combined Ink4ab and Tp53 deficiency revealed a reciprocal senescence and apoptosis regulation, controlling tumorigenesis, including retinoblastoma. INK4-hematopoietic-deficient mice exhibited p19^Arf^-dependent enhanced senescence-like phenotypes, uncontrolled cell proliferation, defective stem cell differentiation, and splenomegaly, with single-splenocytes spatially-enriched in senescence-associated secretory profiles. Our studies reveal the evolutionary origin of paradigms co-regulating senescence and tumor suppression and offer strategies to exploit these reciprocal pathways for cancer prevention and therapy.

## Main Text

The inhibitor of cyclin-dependent kinase 4 (INK4) genomic locus on human chromosome 9p21, and the corresponding locus on mouse chromosome 4, have been subjects to dynamic evolutionary pressure (*1, 2*). In mammals, this small (<50KB) INK4 locus has the highly unusual and remarkable capacity to encode structurally and functionally three distinct cell cycle regulatory proteins: two cyclin-dependent kinase inhibitors, p16^INK4A^ (CDKN2A) and p15^INK4B^ (CDKN2B) (*1*), and a third unrelated protein, p14^ARF^ (p19^Arf^ in mice) translated from an alternative reading frame (ARF) to that of p16^INK4A^ (*3*). These INK4 products share biological properties in regulating cellular senescence and tumor suppression (*2, 4*).

Cellular senescence is a cell phenotype characterized by a durable exit from the cell cycle (*5, 6*). Senescence is initiated in response to nutrient deprivation, mitochondrial dysfunction, oxidative stress, and oncogenic activation (*5*), and comprises cell-autonomous and non-autonomous phenotypes (*7*). The cell-autonomous phenotypes include the acquisition of a flattened morphology, increased reactive oxygen species (ROS), and lysosomal lipid and protein damage, which is marked with lysosomal senescence-associated β-galactosidase (SA-β-gal) (*7*). The cell-autonomous signals are driven by the transcriptional accumulation of p21^Waf1/Cip1^ (CDKN1A), increased INK4 products p16^Ink4A^ and p15^Ink4B^, increased active hypophosphorylated RB1 (Retinoblastoma-1) (*2*), the formation of heterochromatin protein 1 (HP1)- and histone H3 trimethylation at lysine 9 (H3K9me3)-rich chromatin foci, termed senescence-associated heterochromatic foci (SAHF) (*8*), and the associated loss of nuclear lamins, which is p53 or p16^Ink4A^-dependent (*9*). All these cell-autonomous signals culminate to halt cell-cycle progression. The cell-non-autonomous phenotypes are secretory, comprising the senescent-associated secretory phenotype (SASP) (*10*). To accurately identify senescent cells, multiple associated signals and cell features must co-localize (*5*). Damage or stress induces senescence, perhaps to recruit immune cells including macrophages to clear senescent cells (*11*), or elicit an autocrine signaling mechanisms, to maintain the senescent state, suggesting that programmed cell senescence might contribute to physiological functions (*6*). Indeed, programmed cell senescence was found to play instructive roles in tissue remodeling (*11, 12*) and regeneration (*13*). The INK4 product p14^ARF^ (p19^Arf^ in mice) stabilizes p53, a trigger of both apoptosis and senescence, by inhibiting the ubiquitin ligase MDM2 that targets p53 for degradation. ARF is suppressed in some cancers (*14*), and p21^Waf1/Cip1^, a driver of cell cycle arrest in senescent cells, is a key target of p53 transcriptional activity (*12*). Moreover, the p16^Ink4A^/p19^Arf^ locus is repressed by the polycomb repressor complex-1 (PCR1) member and oncogene BMI1, which regulates cellular self-renewal (*15*) and contributes to reduced p21^Waf1/Cip1^ expression (*16*). Notably, in p21-knockout animals, embryonic senescence is partially replaced by a compensatory apoptosis (*11, 12*). However, prolonged cell cycle arrest leads to the upregulation of p16^Ink4A^ (*17*). Single p16^high^ cells were detected in most mouse tissues (*17*) and modulating p16^Ink4A^ levels improves immunoscenescence surveillance (*18*). Additionally, prolonged p16^Ink4a^ activation results in a permanent cell cycle arrest, suggesting that p16^Ink4a^ and possibly other INK4 proteins might orchestrate a reciprocal senescence or apoptotic cell fate.

Programmed cell senescence could be an ancestral characteristic of vertebrates, since senescent cells were detected after fin amputation (*19*), and induced the reprogramming of neighboring somatic cells into stem cells to drive tissue regeneration (*20*). The regenerative effects that senescent cells exert over stem and progenitor cells would likely enhance the risks of tumor formation and therefore have to be controlled by a fail-safe program of tumor suppression. As such, in mammals, p16^Ink4a^ loss results in aging or tumor development (*3, 21*). Here, using comparative functional genomics, we provide evidence for the ancient origin and conserved roles of the INK4 products in senescence regulation and tumor suppression during the vertebrate lifetime. Using retroviral tagging (*22*), we identified an ortholog of INK4 in the zebrafish (Danio rerio) and named *ink4ab*. Using Ink4ab and Tp53 mutant animals, we assessed responses to oxidative and oncogenic stress and the impact of Ink4ab and Tp53 loss on the lifetime tumor incidence. We show that the zebrafish *ink4ab* shares functional attributes with its murine counterpart. Studies in human cells, mouse embryonic fibroblasts (MEFs) and conditional mouse models at the single cell level further elucidate the evolutionary conserved reciprocal cell senescence and apoptotic pathways regulating cell fate through senescence-associated transcriptional and secretory profiles.

## Results

### The evolutionary organization of the INK4 locus

p16^Ink4a^ is the prototype of the family of INK4 proteins, each comprising between 3 and 5 ankyrin repeats (*2*). The ankyrin consensus sequences, which are required for the interactions between p15^Ink4b^/p16^Ink4a^ and CDK4/6 proteins, have been described (*2*). We used these consensus sequences to explore the evolutionary origins of INK4 proteins. Using UniPort, we identified a single gene as a significant homologue of both p15^Ink4b^ and p16^Ink4a^ from cyclostome (lamprey and hagfish) to amphibians (western clawed frog) (Fig. 1). Furthermore, three proteins were identified in *Danio rerio* that show homology to the family of cyclin dependent kinase inhibitors. One locus (ENSDARG00000037262) located on Chromosome 1: 26,127,818-26,131,088 in the reverse strand (GRCz11: CM002885.2) was significantly homologous to both p15^Ink4b^ and p16^Ink4a^ (Fig. 1). Phylogenic analyses indicated that this gene we named ink4ab could be a descendent of ENSGT00940000159801.A, a conserved Vertebrata gene with Vertebrata root species (∼550 million years) among the oldest homologs (Fig. 1, A and B, and fig. S1 to S2).

**Fig. 1.**
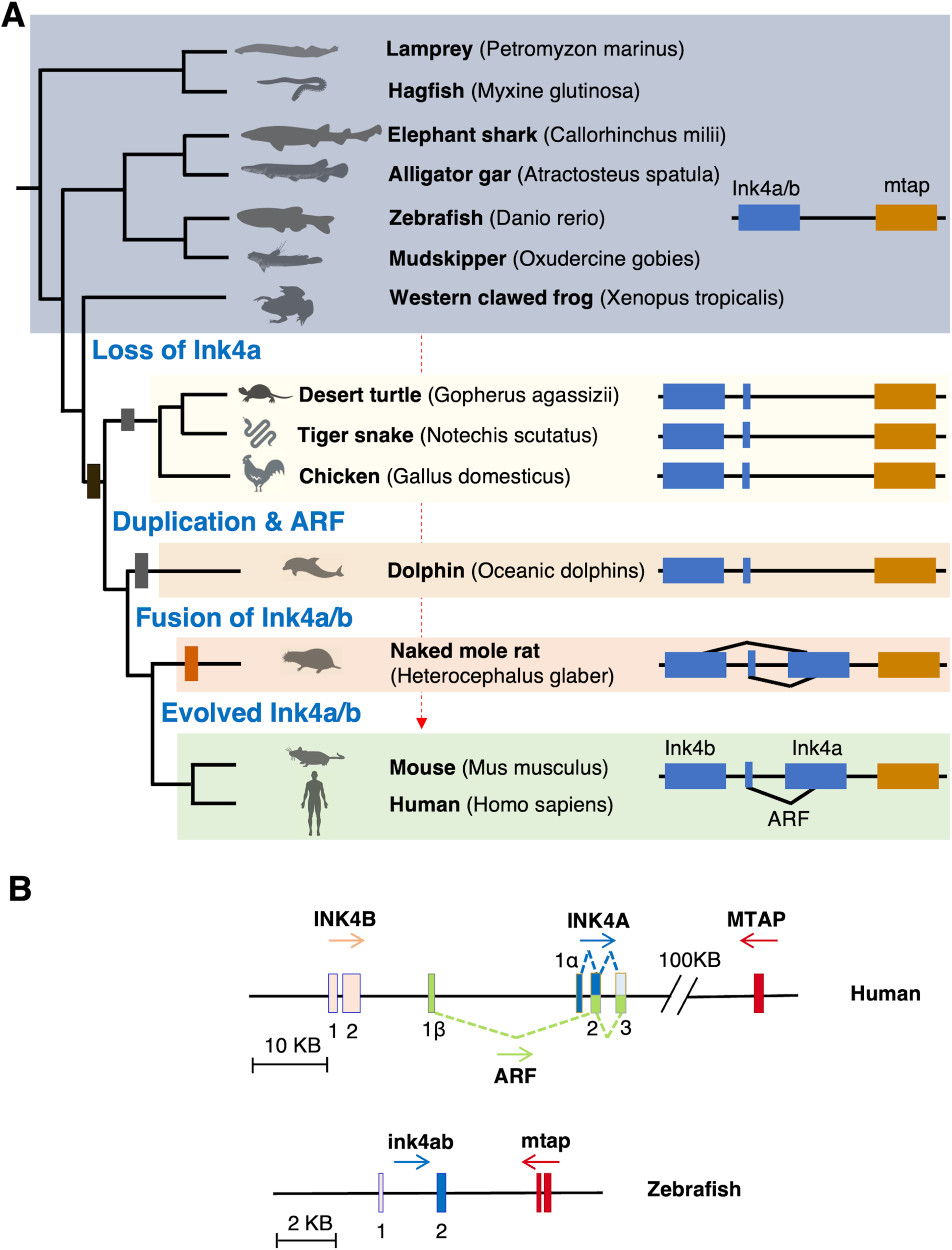
Synteny and phylogenic analysis of zebrafish Ink4ab. (**A)** Proposed evolution of the human and zebrafish Ink4 loci suggesting that the human INK4 genes resulted from a duplication and that the zebrafish INK4 locus contains a single ink4ab gene and no apparent ARF-like sequences. (**B**) The human INK4 locus on chromosome 9p21 is composed of 4 exons (E) – 1α, 1β, 2, and 3 – and encodes two tumor suppressors, p16^INK4A^ and p14^ARF^ (termed p19^Arf^ in the mouse), via alternative reading frames. p16^INK4A^ is translated from the splicing product of E1α, E2, and E3, while p14^ARF^ is translated from the splicing product of E1β, E2, and E3. The CDK inhibitor p15^INK4B^ lies upstream of INK4A and represents the third tumor suppressor gene in this locus. The discovered zebrafish ink4ab is a single gene composed of two exons and lies upstream of the zebrafish mtap gene on Chromosome 1: 26,127,818-26,131,088 in the reverse strand (GRCz11: CM002885.2). Scale is 2 KB.

The genomic structure of this conserved *ink4ab* gene among living jawed fish (chondrichthyans and actinopterygians) is relatively more similar to human p15^INK4B^ than p16^INK4a^. This ancestral *ink4ab* gene consists of two exons and one intron. Syntenic conservation of the genomic regions harboring human p15^INK4b^/p16^INK4a^ and jawed fish *ink4ab* was conserved; the methylthioadenosine phosphorylase (MTAP) gene is adjacent to the human INK4B as in the jawed fish (Fig. 1, A and B). Despite this synteny conservation, a second *ink4* gene was not identified directly upstream in jawless and jawed fish as observed in the human INK4A/ARF locus. In zebrafish, there is no apparent splice variant to indicate the presence of an ARF-like ortholog (fig. S3). We found the zebrafish *ink4ab* not to encode for an equivalent of p16^INK4a^, rather *ink4ab* is a single ancestral gene to both human p15^INK4b^/p16^INK4a^, suggesting that a duplication event occurred after the origin of terrestrial vertebrates likely creating a mammalian locus with the two INK4A and INK4B genes (Fig. 1A and fig. S1 to S2). Moreover, the reading frame for the zebrafish ink4ab does not extend into exon 2, because splicing occurs in a different register to that used in mammals. Therefore, the zebrafish genome has a single p15^Ink4b^/p16^Ink4a^ - like gene and does not appear to contain an ARF (Fig. 1B and fig. S4 to S5).

To determine the expression pattern of *ink4ab* during development of actinopterygians, we decided to use zebrafish, a genetically amiable model organism. Zebrafish *ink4ab* cDNA was synthesized from mRNA isolated from embryos and used to generate a digoxigenin (DIG)-labeled antisense probe to detect ink4ab expression by *in situ* hybridization (ISH). Ink4ab expression was widespread, similar to its mammalian counterparts (fig. S6, A to B), with ISH not revealing any spatially restricted pattern of expression. In the adult wild type (WT) fish, ink4ab mRNA expression was detectable in most tissues examined (fig. S6B), and while levels were low in the heart, kidney, and muscle tissues, ink4ab was strongly expressed in the spleen of adult fish (fig. S6B), suggesting an important role for *ink4ab* in regulating splenocytes.

To study *ink4ab* functions, we designed splicing morpholino oligos (sMO) that interfere with splicing (*23*) and used them to inhibit splicing of exon 2 of *ink4ab*. When injected into the zebrafish embryos at the single cell stage, sMO resulted in *ink4ab* knockdown (fig. S6, C to D). Next, we utilized retroviral insertional mutagenesis (*24*), which allowed the identification of a proviral insertion of the viral long terminal repeat sequences (LTRs) into the *ink4ab* genomic region in fish germ lines (fig. S6C and fig. S7). Carriers for the viral insertion mutation were identified and used for breeding to generate homozygous ink4ab mutants (fig. S7), generating a zebrafish line with an ink4ab insertional mutation in the *ink4ab* upstream regulatory region, frequently regulating gene expression. We confirmed the reduced *ink4ab* expression in heterozygote ink4ab mutant zebrafish and near complete loss of *ink4ab* expression in the ink4ab homozygous mutants (fig. S6D).

### Oxidative and oncogenic stress reveal ink4ab-mediated senescence and tumor suppression

Mammalian p15^INK4b^/p16^INK4a^ are cell cycle regulators that arrest cells at the G1 phase and activate senescence in response to oxidative stress, DNA damage or oncogenic stimuli like Ras activation (*4, 5, 25*). Therefore, we analyzed the cell cycle of ink4ab deficient embryos. As expected, ink4ab deficient embryos have a modest but significant reduction in the proportion of cells in the G1 phase but otherwise similar cell cycle profiles as WT embryos under basal conditions (fig. S6, E to F). Thus, ink4ab deficiency may only be detrimental when cells are under stress that would typically activate senescence. To determine the effect of ink4ab deficiency on the zebrafish response to oxidative stress, WT, *ink4ab* mutants, and *ink4ab* morpholino-injected embryos were subjected to oxidative stress. Senescence was activated in WT embryos, however, *ink4ab* mutant and sMO-injected embryos demonstrated reduced senescence activation in response to oxidative stress (Fig. 2, A and B). RNA sequencing of dissociated cells from embryo proper subjected to oxidative stress revealed highly enriched transcripts, with top transcript p21, followed by ink4ab, among other senescence-related transcripts (fig. S8 to S9A). Ink4ab mutants had higher levels of Atf4 and Rb1 (fig. S9B). We further assessed Suv39hI expression since it is involved in SAHF and transcriptional silencing during senescence (*8*). In WT embryos subjected to oxidative stress, Suv39hI expression increased. However, *ink4ab mutant* embryos had relatively lower Suv39hI expression with and without oxidative stress (fig. S9B). Moreover, while Rb1 expression increased in response to oxidative stress in WT embryos, likely contributing to E2f repression and inducing a cell cycle arrest, Rb1 expression was barely detectable in ink4ab mutants, suggesting a role in the failure to induce senescence during oxidative stress (Fig. 2, A to C).

**Fig. 2.**
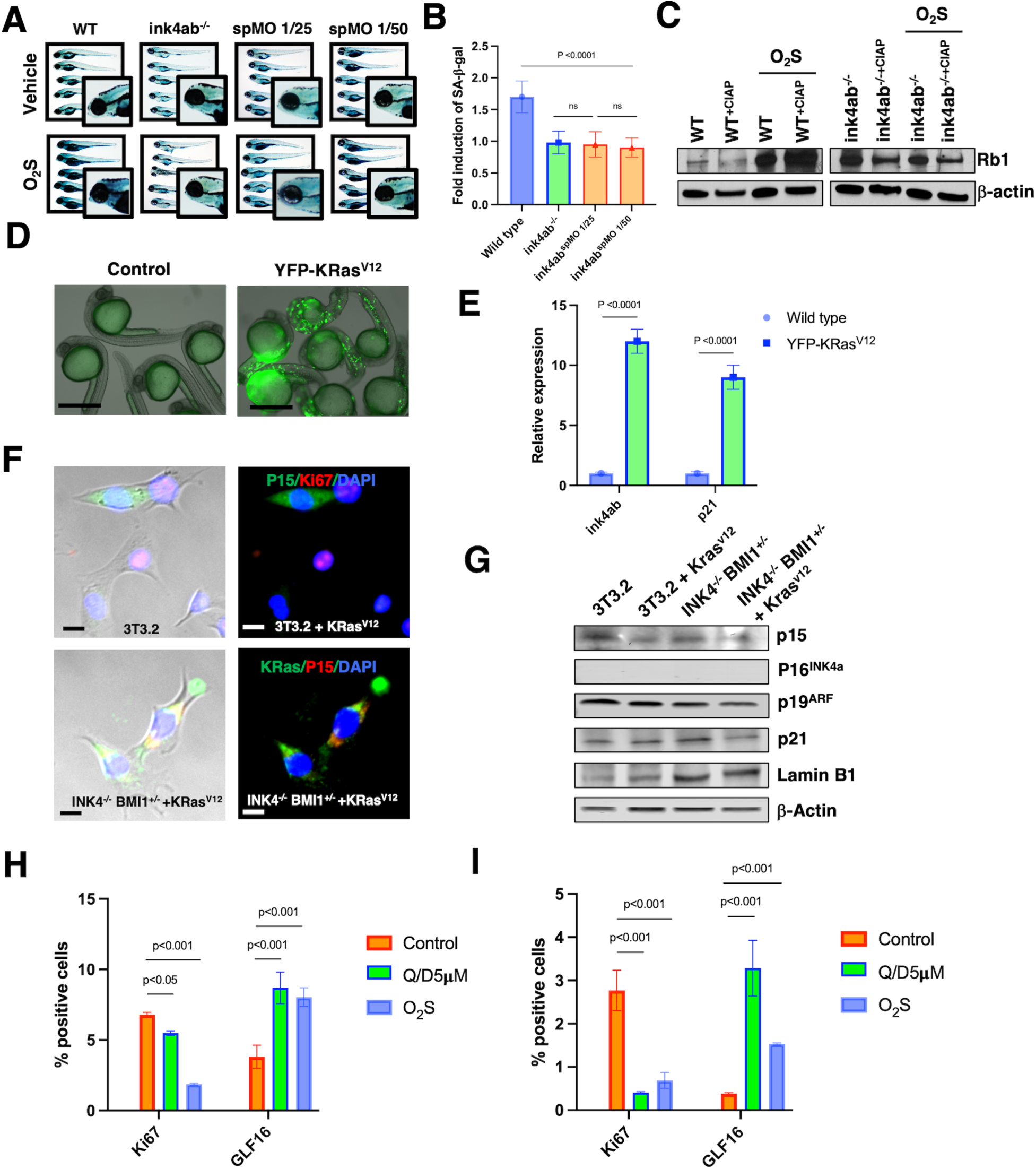
Induction of oxidative and/or oncogenic induced senescence is impaired with INK4 loss. (**A**) Oxidative stress (O_2_S) following H_2_O_2_ treatment resulted in an increase in senescence in wildtype (WT) embryos, but not in mutant or spMO treated embryos with ink4ab deficiency. Senescence-associated β-galactosidase (SA-β-Gal) stained embryos. Inset represents SA-β-Gal densely stained head region in WT but not mutant or spMO treated embryos. (**B**) The average intensity density was used to measure SA-β-gal staining using ImageJ. Data from three independent experiments in which 4-6 embryos were analyzed per sample. (**C**) Western blot confirming decreasing Rb1 expression with ink4ab deficiency in embryos subjected to O_2_S. Lanes with lysates treated with calf intestinal alkaline phosphatase (CIAP) were used to discriminate phosphorylated Rb1 fractions from total Rb1 in untreated lysates. (**D**) Oncogenic Ras expression increases ink4ab expression. Representative images of control vehicle injected embryos and embryos injected with pCMV-YFP-Kras^v12^. (**E**) Oncogenic stress with Kras^v12^ expression resulted in a significant induction of ink4ab and p21 Kras-injected embryos. (**F**) Mouse embryonic fibroblasts with 3T3.2 clone with restored anchorage-independence, isogenic 3T3.2 cells with Kras^v12^ overexpression, Ink4^-/-^/BMI1^+/-^ MEFs with Kras^v12^ overexpression were used to assess senescence induction by p15^Ink4b^. Bright field and IF images of Kras^v12^ overexpression associated with detection of proliferation using Ki67 and p15^Ink4b^ expression. (**G**) In 3T3.2 and Ink4^-/-^/BMI1^+/-^ MEFs that lack p16^Ink4a^, Kras^v12^ overexpression leads to reduced p15^Ink4b^ and activation of the senescence marker p21. A reduction in Lamin B1 was not clearly notable under these experimental conditions. Data from six independent experiments. (**H**) Quantitation of the levels of proliferating cells by Ki67 positive cells and GLF16-senescence-associated β-Gal positive cells in transformed mouse CT26 colorectal cancer (CRC) cells, which are deficient in p15^Ink4b^/p16^Ink4a^ but have p53^WT^ (*29*). Note that senescence induction and/or oxidative stress reduced Ki67 levels and increased GLF16-senescence-associated β-Gal. (**I**) In MC38 cells, which are CRC cells deficient in p15^Ink4b^/p16^Ink4a^ and harbor a mutant p53 (*29*), less senescence inductions could be achieved. Note the much lower basal GLF16-senescence-associated β-Gal levels relative to CT26 cells. Data in **H** and **I** represent six independent experiments. Scale bars are 100 μM in **D** and 20 μM in **F**.

We hypothesized that loss of *ink4ab*, and therefore less senescence activation, could signal Tp53 to increase apoptosis as a mechanism to prevent carcinogenesis. To test this hypothesis, we treated embryos with campothecin (CPT), a topoisomerase inhibitor, which induces DNA breaks and activate apoptosis. Ink4ab mutants demonstrated strikingly higher levels of apoptosis with CPT treatment but also displayed significantly higher basal levels of apoptotic cells (fig. S9, C to D). Knockdown of Tp53 with MO dramatically reduced the number of apoptotic cells in both untreated and CPT-treated ink4ab mutant embryos (fig. S9, C to D). This suggests that ink4ab mutants have increased activation of Tp53-dependent apoptosis. Altogether, these data suggest a key role of ink4ab in cell cycle signaling during oxidative stress.

The ability of Ras oncogenes to engage tumor suppressor pathways limits tumor cell outgrowth and activates senescence and/or cell death (*25*). The outcomes could be oncogene-induced senescence (OIS) and oncogene-induced apoptosis (OIA), two crucial tumor suppressor checkpoints used to restrain tumor growth (*4*). We hypothesized that *ink4ab* may play a role in activating OIS. To test this hypothesis, the Kras-YFP mRNA was injected into single cell embryos to induce oncogenic stress (Fig. 2D). Oncogenic Ras expression resulted in a significant increase in not only ink4ab (20-fold), but also p21 (10-fold) (Fig. 2E). However, as in mammals (*4*), Ras overexpression resulted in embryonic death in zebrafish, limiting the long-term assessment of Ras effects in the adult zebrafish.

We next performed a cross-species study of *ink4ab* to determine if the role of *ink4ab* in responding to oncogenic stress is evolutionarily conserved. We synthesized and expressed an inducible *ink4ab*-EGFP fusion (to detect the fusion protein using an EGFP antibody) or mouse p15^Ink4b^ (mp15) into MEFs, which lack p15^Ink4b^/p16^Ink4a^ through a homozygous deletion (*26*) and assessed responses to Ras activation. Similar to mp15, the zebrafish Ink4ab reduced the proliferation of Hras^V12^ expressing cells (fig. S10, A to B). Since p15^Ink4B^ expression should modulate RB1 phosphorylation through inhibition of cyclin D/CDK4/6 active complex formation, we detected changes in pRB^Ser780^ phosphorylation with both mp15 and zebrafish Ink4ab, as expected from a functional conservation (fig. S10, B and D). We similarly examined the effects of *ink4ab* in human 293T cells in combination with HRas^V12^ for cell cycle changes using flow cytometry. Similar to results of pRB phosphorylation changes under basal conditions, we found no difference in the cell cycle profiles under basal conditions. However, upon oncogenic stimulation, there was a modest increase in the percentage of cells in G1 and a decrease in the percentage of cells in S phase of the cell cycle observed in cells expressing both ink4ab and oncogenic HRas (fig. S10E).

The INK4 locus is repressed by the polycomb repressor complex (PRC1) member and oncogene BMI1 (*16*). PRC1 proteins are transcriptional repressors causing chromatin compaction, which allows access to the INK4 locus to facilitate senescence (*16*). Moreover, cells cease proliferation in culture when they are spatially confined by their neighbors, a phenomenon called contact inhibition. During contact inhibition, cells do not undergo senescence (*27*). Immortalized MEFs lose their strong contact inhibition over time. We utilized a new cell clone (3T3.2) derived by single cell cloning with restored contact inhibition to study senescence in response to oncogenic Ras activation. Kras^V12^ activation in either 3T3.2 cells or p16^Ink4A-/-^/BMI1^+/-^ MEFs resulted in increased senescence-associated β-gal, which was detected upon optimizing the use of a fluorescent substrate (fig. S11, A to C) and validated with a fluorescent small molecule (GLF16) developed to detect senescent cells (*28*) (fig. S11, D to F). Notably, increased senescence-associated β-gal was associated with reduced p15^Ink4B^ positive cells (Fig. 2, F and G, and fig. S12, A to C), increased cell proliferation, and increased Ki67 positive cells in the case of p16^Ink4A^/BMI1^+/-^ cells (fig. S12, D to F). We next assessed the senescence levels in already transformed mouse colon cancer cells. In CT26 cells which are deficient in p15^Ink4b^/p16^Ink4a^ but are p53^WT^ (*29*), senescence induction and/or oxidative stress reduced Ki67 levels and increased GLF16-senescence-associated β-Gal (Fig. 2, H, and fig. S13A), while in MC38 cells which are deficient in p15^Ink4b^/p16^Ink4a^ and harbor a mutant p53 (*29*), less senescence inductions could be achieved, and moreover, the basal GLF16-senescence-associated β-Gal levels were much less (Fig. 2, I, and fig. S13, B to D). These data suggest a reciprocal regulation in senescence induction between INK4 products, revealing that with p15^Ink4b^/p16^Ink4a^ loss, p19^Arf^ could compensate, and when p19^Arf^ is also lost, p53 regulates the senescence-associated responses to oxidative and/or oncogenic stress.

### The tumor suppressor functions of ink4ab in zebrafish

A cohort of zebrafish (n=312) was analyzed for survival over their average lifespan (Fig. 3A). The mean survival for WT (n= 124) and ink4ab^+/-^ fish (n= 27) was 22.44±0.21 months and 18.1±0.92 months, respectively (table S1). The survival of the ink4ab^-/-^ fish (n= 161) was 16.7±0.5 months and was significantly lower in pairwise comparisons (p<0.001, log-rank test and Sidak multiple comparison). A Cox proportional hazard model using WT fish as a reference group showed that ink4ab^+/-^ fish has a 258% higher risk of death, and ink4ab*^-/-^* fish had a 764% higher risk of death, respectively, than WT (table S1). For the ink4b^+/-^ fish, the risk of death is 59% lower (Hazard ratio=0.41) than ink4ab^-/-^ fish. Some aspects of aging have been argued to benefit from the tumor suppressor functions of mediators of senescence such as p53, ARF, and p16^INK4a^ (*30*). Since ink4ab fish showed lower survival, they were frequently identified with an extreme pallor and often with focal melanocytic pigmentations (Fig. 3B). Senescence increases with age in zebrafish and can be detected using β-gal staining (*31*). Since we detected the role of *ink4ab* in activating senescence signals during oxidative and oncogenic stress in embryos, we assessed senescence levels in older adult fish. The ink4ab deficient adult zebrafish had a significantly lower senescence-associated β-gal relative to WT (Fig. 3, C and D). As CDKN2A (Ink4A/Arf) knockout mice had enlarged spleens, which are indicative of abnormal hematopoiesis (*3*), and since splenomegaly is a common manifestation of splenocyte hyperproliferation, we examined dissected spleens from cohorts of WT and ink4ab^-/-^adult fish (n=5/group) at 8 months of age. Splenomegaly was observed in all ink4ab mutants analyzed, with abnormal blast-like cells and enlarged spleens, to a size of almost two times larger than that of WT animals (Fig. 3, E to G).

**Fig. 3.**
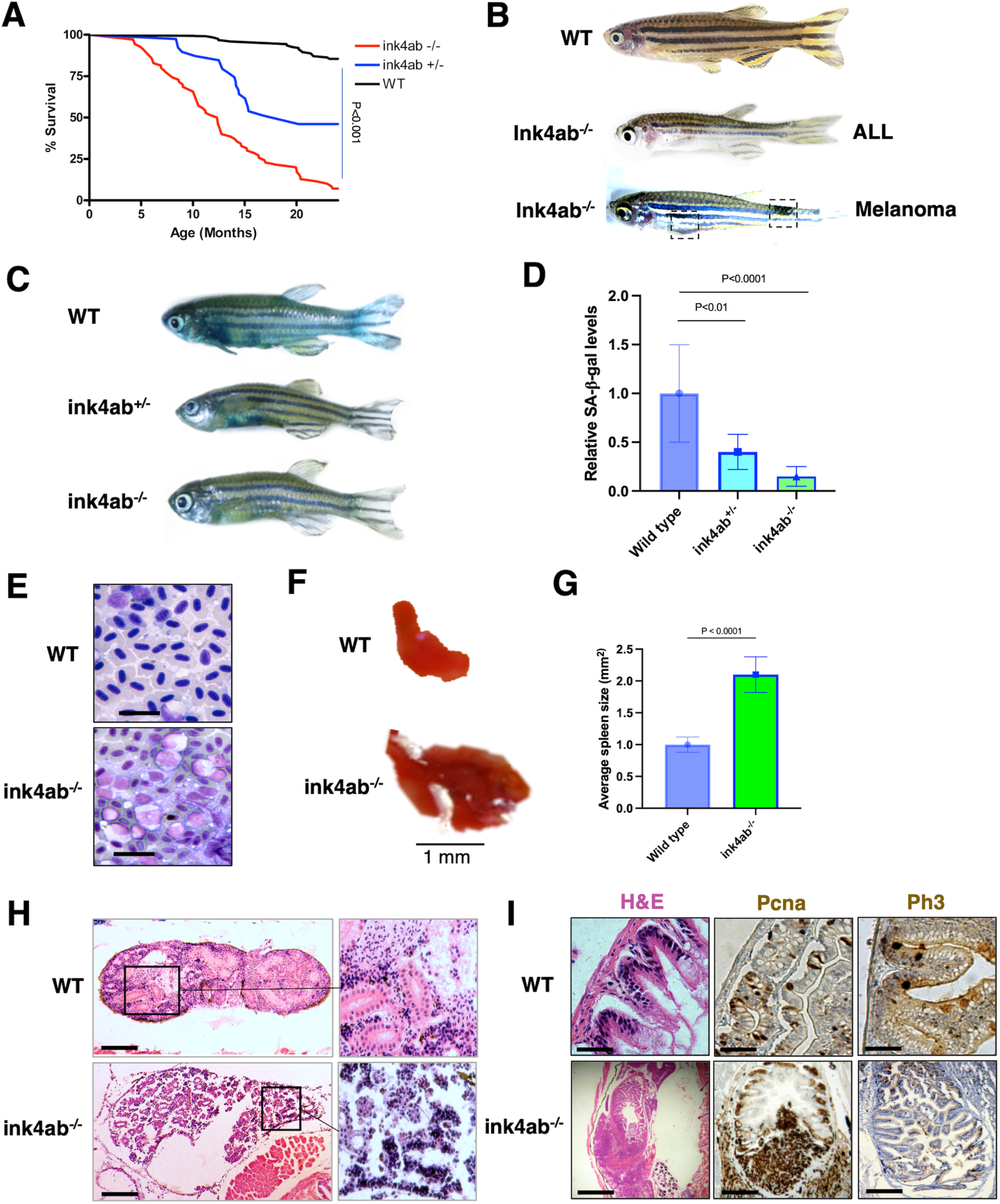
Phenotypes of zebrafish with ink4ab deficiency. (**A**) Ink4ab deficiency decreases fish survival. An average lifespan 2-year Kaplan-Meier survival curve of WT and ink4ab mutant zebrafish. (**B**) Representative images of ink4ab adult fish with melanoma and ALL phenotypes. Note the abnormal melanocytic pigmentations in the tail, fin, and abdominal regions. (**C**) Zebrafish *ink4ab* deficiency correlates with a decrease in age-related senescence. Whole mount SA-β-Gal staining in adult zebrafish. (**D**) Quantitation of SA-β-Gal staining in the adult fish with ink4ab mutations. (**E**) A touch prep of spleen cells stained with Giemsa showing increased number of lymphoid and blast-like cells in the enlarged ink4ab splenocytes relative to WT spleens. (**F**) Splenomegaly observed in ink4ab mutants. (**G**) Representative spleens dissected from WT and ink4ab mutant fish and measured along the longitudinal axis to display spleen sizes. (**H**) Pancytopenia with lymphoid-like leukemia in the kidney marrow of a 12-month-old ink4ab mutant zebrafish. These fish were analyzed for blood cancers by obtaining peripheral blood smears from clipping the fin or extracting blood from the heart as well as analyzing whole kidney marrow sections stained with H&E. (**I**) Intestinal cancer phenotype in a 14-month-old ink4ab mutant fish with an abdominal swelling. Top images demonstrate WT intestine and bottom images of ink4ab mutant intestine with hyperproliferation and disorganization of intestinal crypts, suggestive of intestinal cancer. The H&E phenotype was confirmed with IHC staining for Pcna (demonstrating hyperproliferation) and Ph3 for cell cycle phenotype. Scale bars are 20 μM in **E** and 100 μM in **H** and **I**.

A cohort of ink4ab mutant fish was monitored for spontaneous tumor development. Several tumor types were identified, including leukemias and solid tumors. Lymphocytic leukemias were observed at an incidence of 4% in the ink4ab^-/-^ fish (Fig. 3H). Other tumor types were typically only revealed after histological analyses, including metastatic melanoma, osteosarcoma, intestinal cancer, and hepatocellular adenoma (Fig. 3I). Melanocytic lesions in the ink4ab^-/-^ fish were metastatic in two cases and metastasis was detected using a melanocyte-specific Schmorl’s stain (fig. S14 A to C). As p16^INK4A^ loss is frequent in hereditary human melanomas and somatic p16^INK4A^ loss accelerates melanomagenesis (*32*), it was no surprise to observe melanomas develop in ink4ab mutants (Fig. 3B and fig. S14). To study the effects of ink4ab loss on the hematopoietic recovery and response to DNA damage upon γ-irradiation, we treated cohorts of a year-old WT and ink4ab^-/-^ zebrafish with a sublethal myeloablative dose of radiation (*33*) (23 gray, Gy) and then monitored them for survival. The ink4ab^-/-^ fish subjected to irradiation (n=12) had a mean survival of 31.5 days and all died by day 52 post-radiation compared to only one death from the cohort of matched irradiated WT fish (n=12) (fig. S15A). The overall survival of irradiated ink4ab^-/-^ fish was significantly less (p< 0.001, log-rank test) compared to irradiated WT fish (table S2), with ink4ab^-/-^ fish having a 1,528% higher risk of dying from myeloablative radiation than fish with no radiation (HR=16.28).

### Combined ink4ab and p53 loss in the lifetime of zebrafish

We next crossed the p53 (tp53^M214K^) mutant zebrafish (*34*) to ink4ab mutants to develop a line deficient for both tumor suppressors. We did not detect tumors in the cohort of zebrafish heterozygous for tp53^M214K^ mutation alone by 16 months of age, consistent with prior studies (*34*). However, zebrafish heterozygous for both tp53^M214K^ and ink4ab mutation displayed a lower incidence of malignant tumors (22.5%) comparable to the incidence observed in homozygous tp53^M214K^ mutants (28.0%) (table S3), suggesting a potential synergistic tumor suppressor activity between p53 and ink4ab. Furthermore, the ink4ab mutation status influenced the age of tumor onset, tumor number, and tumor type (Fig. 4, A to C and table S3). By 16 months, 75.0% of homozygous mutants for both p53 and ink4ab had developed various tumor types, including nerve sheath tumors (37.5%), retinal tumors (31.3%), and others (6.3%, majority melanomas) (Fig. 4C and table S3). A logistic regression model was used to determine the association of tumor incidence with different genotypes involving *ink4ab* and p53 mutations. p53^+/-^ was used as a reference group (fig. S15B). Moreover, the odds of developing a tumor for a fish heterozygous for p53 mutation was 97.5% lower than the odds for fish with p53^-/-^ ink4ab^-/-^ (fig. S15B). Interestingly, ink4ab^-/-^ fish had more and larger variety of tumor types than the p53^+/-^/ink4ab^+/-^ fish (Fig. 4C), while the homozygous loss of both ink4ab and p53 in the p53^-/-^/ink4ab^-/-^ was detrimental with majority loss of tumor suppressor functions (Fig. 4C and fig. S15B).

**Fig. 4.**
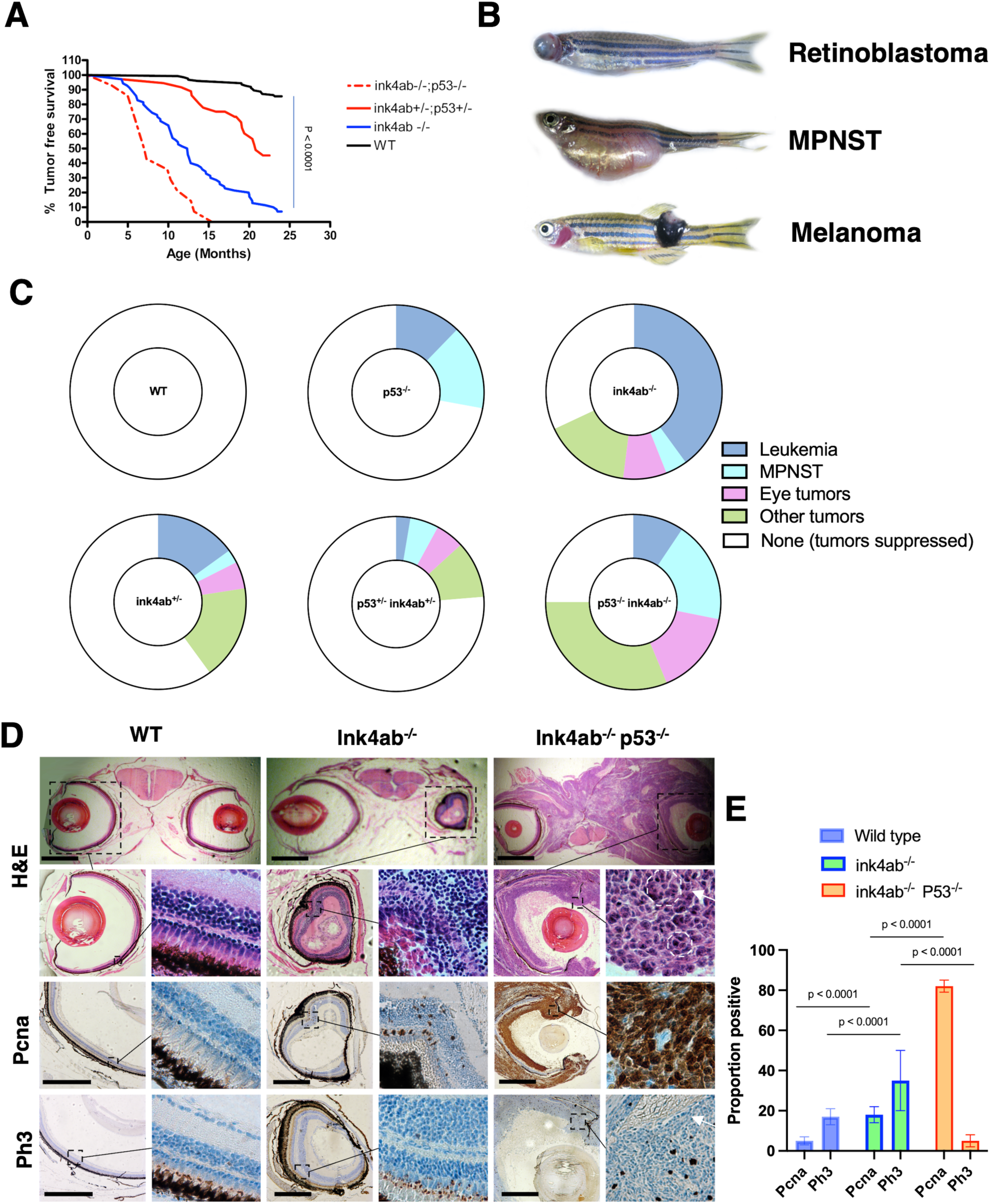
Zebrafish doubly deficient for ink4ab and p53 have reduced tumor-free survival. **(A**) Kaplan-Meier survival curve for ink4ab/p53 mutants. (**B**) Tumors frequently observed in ink4ab/p53 double mutant fish included retinoblastoma, malignant peripheral neural sheath tumors (MPNST), and occasionally melanomas. (**C**) Proportion of zebrafish with tumors suppressed correlates with allelic loss of ink4ab or p53, suggesting synergistic tumor suppressor activity between p53 and ink4ab, where mono-allelic loss of either or both inkab and p53 is associated with more tumor suppression than bi-allelic loss of either. Colors indicate the tumor types while in WT (White), there was complete tumor suppression. (**D**) Combined loss of ink4ab and p53 increases retinoblastoma tumor progression. WT gross histology and H&E sections of retina. ink4ab^-/-^ mutant fish frequently observed with unilateral eye deformity, as shown with left eye deformity associated with retinal layer invasion. The combined ink4ab/p53 double mutant fish demonstrated an aggressive bilateral eye tumor with retinoblastoma like features. Retinal invasion and features of retinoblastoma such as Flexner-Wintersteiner rosette (white outline and arrow) are seen. These rosettes (named for pathologist Simon Flexner and ophthalmologist Hugo Wintersteiner) consist of tumor cells surrounding a central lumen that contains cytoplasmic extensions from the tumor cells. (**E**) Levels of Pcna and Ph3 in single ink4ab and double ink4ab/p53 mutant fish retinal layers. Images on the right are 10X magnifications of the outlined areas within the images on the left. Scale bars are 500 μM.

Retinal tumors represented unique and readily visible aspects of ink4ab loss (Fig. 4B and Fig. 4D). Analysis of these eye tumors revealed disruption of the retinal layer and retinoblastoma-like features, such as Flexner-Wintersteiner rosettes of primitive photoreceptor cells (*35*) detected upon histological examination, which are commonly used to diagnose retinoblastoma in humans (Fig. 4D). These tumors were present in ink4ab^-/-^ and appeared bilateral, more aggressively larger, faster growing, and metastatic into the surrounding tissues including the brain in the p53^-/-^/ink4ab^-/-^ fish (Fig. 4D), with notably a high percentage of proliferative Pcna and mitotic Ph3 positively stained cells (Fig. 4E). Together, these findings demonstrate the tumor suppressor functions of ink4ab and enhancement of p53-mediated tumor suppressor effects in zebrafish, as seen in mammals.

### Conditional compound INK4 products’ deficiency in mouse HSCs is tumorigenic

Frequently in human cancers including leukemias, 9p21 deletions interfere with all INK4 proteins thus also remove p19^Arf^ and modulate p53 functions. In mouse models, the effects of combined p16^Ink4A^ and p15^Ink4b^ loss have been examined (*36*), however, the role p19^Arf^ in hematopoietic cells under conditions of p16^INK4A^ and p15^Ink4b^ loss remains ambiguous. Therefore, we examined the effects of the conditional deficiency p19^Arf^ in the mouse hematopoietic stem and progenitor cells (HSCs) when combined with p16^Ink4A^ and p15^Ink4b^ deficiency. We utilized the Vav-Cre male mice crossed with female p15^Ink4b^/p16^Ink4a^ mutant mice with either one wild type (WT) and one floxed p19^Arf^ allele (Ink4ab/Arf^+/-^Vav-Cre^+^) or both p19^Arf^ floxed alleles (Ink4ab/Arf^-/-^Vav-Cre^+^) (Fig. 5A). Cre expression by the vav regulatory elements, which are able to drive hematopoietic tissue-specific cre expression in the thymus, bone marrow, and spleen (*37*) to mediate mono-allelic (Ink4ab/Arf^HSC+/-^) or bi-allelic (Ink4ab/Arf^HSC-/-^) loss of p19^Arf^ expression in HSCs of mice deficient in p16^Ink4A^ and p15^Ink4b^ (Fig. 5, A to B). Mice were born at the predicted mendelian frequency and were normal during development but showed allelic-dependent impaired survival (Fig. 5B). Peripheral blood smears showed an increased number of progenitor cells in the Ink4ab/Arf^HSC+/-^ mice (fig. S16A), while the Ink4ab/Arf^HSC-/-mice^ presented with anemia and leucopenia (fig. S16A). Bone marrow (BM) analyses suggested a myelodysplastic-like phenotype in Ink4ab/Arf^HSC+/-^ mice (fig. S16B), while the Ink4ab/Arf^HSC-/-^ mice revealed BM infiltration with homogenous blast-like cells, suggestive of a leukemia/lymphoma-like phenotype (fig. S16B). Ink4ab/Arf^HSC-/-^ mice analysis showed enlarged organs, and in particular, the spleens were greatly enlarged in size (Fig. 5C and fig. S16C).

**Fig. 5.**
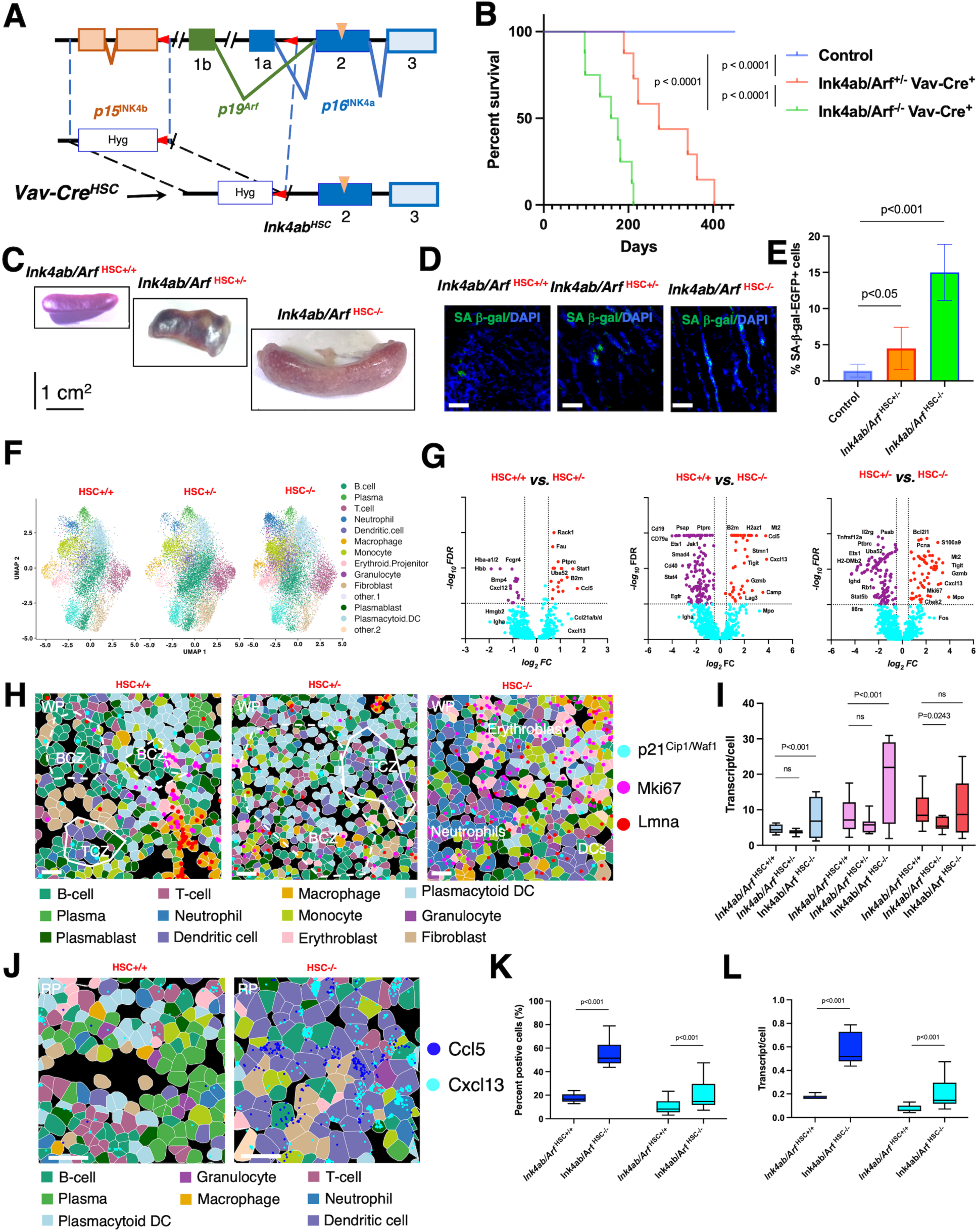
Single cell spatial studies of splenocytes from the conditional INK4ab/Arf mutant mice. (**A**) Experimental design for generating the conditional mouse model of combined p16^Ink4A^, p15^Ink4b^, and p19^Arf^ loss in hematopoietic cells using the Vav-cre mice. Diagram demonstrates mutations in both p16^Ink4A^ and p15^Ink4b^, (inverted triangles), and floxed p19^Arf^ resulting in complete loss of all three Ink4 proteins in the mouse HSCs. (**B**) Kaplan-Maier survival curve of control and cre-recombined Ink4 mice (n =12/group). (**C**) Splenomegaly dosage dependent phenotype in the mono-allelic (Ink4ab/Arf^HSC+/-^) or bi-allelic (Ink4ab/Arf^HSC-/-^) loss of p19^Arf^ expression in HSCs of mice deficient in p16^Ink4A^ and p15^Ink4b^. (**D**) IF images of mouse splenocyte sections stained with senescence-associated β-Gal-EGFP probe. (**E**) Quantitation of senescence-associated β-Gal-EGFP levels. (**F**) Single cell spatial analyses of splenocyte phenotypes. Cellular neighborhoods in CosMx data visualized by Uniform Manifold Approximation and Projection (UMAP) using semi-supervised cell clustering of mouse splenocytes from control and INK4 mutant mice. (**G**) Differential expression using a two-sided Wilcoxon Rank Sum test between splenocytes in Ink4ab/Arf^HSC+/+^ *vs.* Ink4ab/Arf^HSC+/-^, Ink4ab/Arf^HSC+/+^ *vs.* Ink4ab/Arf^HSC-/-,^ and Ink4ab/Arf^HSC+/-^ *vs.* Ink4ab/Arf^HSC-/-^ in the Red pulp and White pulp areas (cells = 26,440 from Ink4ab/Arf^HSC+/+^) compared to other neighborhoods (cells = 12,582 from Ink4ab/Arf^HSC+/-,^ and cells = 35,768 from Ink4ab/Arf^HSC-/-^), with top twelve transcripts enriched or reduced between samples and related to senescence and/or splenocyte proliferation being shown in Ivy-gap data (n = 35 FOVs across 6 samples). (**H**) Representative FOV multichannel overlay of senescence-associated p21^Cip1/Waf1^ (light blue), proliferation signal MKi67 (pink), and nuclear Lamin (red) detected from the 980-plex assay, morphological, and cell typing markers. BCZ, B-cell zone, TCZ, T-cell zone, DCs, dendritic cells, WP, white pulp. Specificity of p21^Cip1/Waf1^ spatial distribution was evaluated by lack of detection of the proliferation marker Ki67 in the same spatially oriented single splenocytes within each cell neighborhood and FOV. See **fig. S18C** for individual senescence-associated markers in the same FOVs used to confirm the specificity of p21^Cip1/Waf1^, Lamin, and Mki67 staining in single splenocytes. A mutually exclusive relationship between p21^Cip1/Waf1^ and Mki67 positivity was depicted. (**I**) Levels of p21^Cip1/Waf1^, Mki67, and nuclear Lamin between splenocytes from control and mutant mice. (**J**) Representative FOV multichannel overlay of Spatial domain distribution of splenic microenvironment clusters with SASP cytokines Ccl5 and Cxcl13 most enriched in cellular neighborhoods of neutrophils and DCs, respectively, and detected from the 980-plex assay and morphological markers. For each cell, the nearest neighbors were identified, and a summary of those neighbors was recorded. The abundance of each cell type is used. This operation is performed for all cells in FOVs, defining a matrix of cells and neighborhood characteristics. (**K**) Increased number of cells positive for Ccl5 and Cxcl13 in splenic neutrophils and DCs of Ink4ab/Arf^HSC-/-^ *vs.* Ink4ab/Arf^HSC+/+^ (n = 6 and 14 FOVs from duplicate samples). (**L**) Ccl5 and Cxcl13 transcripts are enriched in cell neighborhood of neutrophils and DCs of Ink4ab/Arf^HSC-/-^ splenocytes. Scale bars are one cm in **C**, and 100 μM in **D**, **H**, and **J**.

### Enhanced senescence and hypersecretory neighborhoods from single Ink4ab/Arf splenocytes

The spleen functions as the largest secondary lymphoid organ of the body and splenocytes play major roles in hematopoiesis, immune functions, red blood cell clearance, and potentially senescence regulation (*38*). The mouse spleen is structurally divided into the majority tissue red pulp (RP) and less than a quarter of white pulp (WP), separated with the marginal zone (MZ) (*38*). The WP is the primary adaptive immune response region, including T-cells in the T-cell zone (TCZ) and B-cells in the B-cell zone (BCZ) follicles, while the RP contains innate immune cells, including neutrophils, monocytes, dendritic cells (DCs), gamma delta (ψ8) T-cells, and macrophages (*38*). Since the spleens of INK4 mice were greatly enlarged, we first assessed the levels of senescent cells in splenocytes. Senescence β-gal positive cells were significantly increased in the splenocytes of Ink4ab/Arf^HSC+/-^ and Ink4ab/Arf^HSC-/-^ mice (Fig. 5D) and in liver cells of Ink4ab/Arf^HSC-/-^ mice (fig. S17A). Notably, histological examination revealed severe disruption of the splenic architecture in the Ink4ab/Arf ^HSC-/-^ spleens, including loss of the MZ, blurring of the RP-WP distinction, and disrupted TCZ and BCZ follicles (fig. S16C and fig. S17D to S17E).

To identify the effects of Ink4ab/Arf loss in HSC differentiation and senescence regulation, we optimized a new high-plex assay for monitoring single cell spatial profiles of mouse splenocytes using the CosMx Spatial Molecular Imager (SMI) (*39*). We then characterized the single-cell spatial distribution of splenic cell phenotypes with spatially resolved single cell transcriptome analyzed in splenic sections from Ink4ab/Arf^HSC+/+^ (control), Ink4ab/Arf^HSC+/-^ and Ink4ab/Arf^HSC-/-^ mice. A total of 74,790 single-cell transcriptomes were obtained, including 26,440 transcriptomes from controls and 48,350 transcriptomes from Ink4ab/Arf^HSC+/-^ and Ink4ab/Arf^HSC-/-^ mouse spleens (Fig. 5F and table S4). Furthermore, 14 clusters representing distinct cell populations were identified (Fig. 5F and fig. S17C to D). We identified distinct potential cell identities between control and Ink4ab/Arf^HSC-/-^ splenocytes and found differences in the proportion of each type of cell (fig. S17, D to F). The proportion of B-cells, T-cells and NK-cells was decreased, while the proportion of granulocytes and monocytes was increased in the Ink4ab/Arf^HSC-/-^ splenocytes (fig. S17D to E), which suggested that the cells in these clusters could be involved in the senescence response to dysfunction or damage. Thus, we focused on the significant differentially expressed genes (DEGs) of these cells between control and either Ink4ab/Arf^HSC+/-^ or Ink4ab/Arf^HSC-/-^ (Fig. 5G and table S5). We clustered the DEGs from control *vs.* Ink4ab/Arf^HSC+/-^, control *vs.* Ink4ab/Arf^HSC-/-^, and Ink4ab/Arf^HSC+/-^ *vs.* Ink4ab/Arf^HSC+/-^ (Fig. 5G). The results showed enrichment in Rack1, Fau, Ptprc (B220), Stat1, B2m, and Ccl5 in the Ink4ab/Arf^HSC+/-^ splenocytes, and diminished Hbb, Hba-11/2, Fcgr4, and Cxcl12 (Fig. 5G). DEGs between control and Ink4ab/Arf^HSC-/-^ revealed enrichment in the senescence regulator tissue stress alarmin signal Hmgb2 and expression of Cxcl13, Ccl5, H2az1, B2m, Stmn1, Tigit, Gzmb, and lag3, which include immune checkpoint and SASP targets that were further enriched when comparing Ink4ab/Arf^HSC+/-^ *vs.* Ink4ab/Arf^HSC+/-^, and additionally accompanied with enriched proliferation transcripts including Pcna and Mki67, suggestive of a tumorigenic phenotype (Fig. 5G).

Since B-cells play important roles in humoral immunity, they are the largest cell population that we detected in the spleens of control and INK4 mice. In fact, B cells constituted the major cell population (54.7 %) in the control mice and constituted approximately 44.2 % of the cells in the Ink4ab/Arf^HSC+/-^, but they were significantly diminished in the Ink4ab/Arf^HSC-/-^ mice with apparent retention of CD34^+^ cells (Fig. S17C to E). Using custom R scripts, microdomains containing senescence-like cell phenotypes were identified on the basis of validated senescence-related markers with the highest sensitivity and specificity to detect senescent cells *in situ/in vivo* (*5*), and considering that our splenocytes harbor mutations in p15^INK4b^/p16^INK4a^ and conditional loss of p19^Arf^. We utilized p21^Cip1/Waf1^ as a core marker of cellular senescence in cells without cell cycle progression as approximated by lack of Pcna and Mki67 transcripts, and included nuclear envelop lamin component Lmna in the same splenocytes (Fig. 5H and fig. S18 A to C). Splenocytes in the BCZ upregulated p21^Cip1/Waf1,^ while more proliferative Mki67+ splenocytes were enriched in the WP near plasma cells and plasmablasts and Lmna+ splenocytes widely distributed (Fig. 5H and 5J). The BCZ follicles contain follicular DCs (FDCs) which present antigen to follicular B-cells while also producing CXCL13 to help organize the B-cell follicle (*38*). We observed B-cells typically gathered in dense clusters accompanied by T-cells (Fig. 5H). In INK4 splenocytes, plasmablasts gathered densely, often proximal to smaller numbers of T-cells. Macrophages and DCs both gathered in disturbed clusters and trafficked diffusely throughout RP (fig. S18C). Moreover, the Ink4ab/Arf^HSC-/-^ splenocytes were enriched in neutrophils, monocytes, and erythroid progenitors at the expense of B- and T-cell lineages (fig. S18C). This spatial information also allows detailed analysis of cell neighborhoods. We defined a neighborhood matrix encoding the number of each cell type among each cell’s 200 closest neighbors (Fig. 5J). The gene expression levels of CXCL13 and CCL5 SASP cytokines were particularly increased in Ink4ab/Arf^HSC-/-^ cells compared to control cells (Fig. 5K to 5L). In addition, increased senescence-related transcripts was associated with immune-related pathways, DNA damage repair-related pathways, and the activation of several oncogenic induced senescence processes, such as enriched Check1 and stathmin-1, loss of Pten within the DNA damage repair (DDR) and deregulated metabolic senescence subtype and activated components of SASP (Fig. S19A to B). We confirmed the altered B-cell and T-cells differentiation trajectories in these splenocytes (fig S20), and the enhanced proliferation (Pcna, Mki67), upregulated cyclin D, and altered HP1 and H3K9Me forming SAHP (fig. S21). These data suggest that conditional loss of p19^Arf^ when associated with lack p15^Ink4b^/p16^Ink4a^ generates a repertoire of senescence-like phenotypes in the likely pro-tumorigenic environment, leading to an uncontrolled cell proliferation and defective stem cell differentiation, with single-splenocytes spatially enriched in senescence-associated secretory profiles.

## Discussion

We describe the ancient origin of the INK4 products and their functions in senescence regulation and tumor suppression. Using functional genomic elements, in this case ankyrin repeats (*2*), we identified evolutionary-conserved ink4 genes and utilized developmental genetics for modeling cell cycle regulation and studying how INK4 proteins might orchestrate a reciprocal senescence or apoptotic cell fate (*24, 31, 33, 34, 40*). Orthologs of INK4 have been identified in Xiphophorus fish and Fugu (*41*), among other vertebrates and shown to functionally trigger senescence upon oxidative or oncogenic stress. In the naked mole rat, a species with extremely low rates of cancer, the INK4 locus encodes an additional product that consists of p15^Ink4b^ exon 1 joined to p16^Ink4a^ exons 2 and 3 and may be the result of an early translation termination event.

This naked mole rat isoform is absent in human and mouse cells (*42*) and induced during early contact inhibition and senescence (*42*). We found the homology between the naked mole isoform and zebrafish ink4ab to be relatively low, suggesting that INK4 proteins evolved into contingent tumor-suppressive mechanisms that trigger senescence or apoptosis based on sensors and transducers in a coordinated and cell-specific networks. Third, due to its characteristic of limiting cell proliferation, cellular senescence was categorized as a tumor-suppressor mechanism (*25*), however, depending on the context, senescence plays both beneficial and detrimental roles during tumor progression. Our results solidified the tumor-suppressor functions associated with senescence in the presence of p19^Arf^ or its ortholog *ink4ab*, and the tumor-promoting actions of senescent cells via SASP in mouse spleens in its absence. Mice individually lacking p19^Arf^ and p16^Ink4a^ effectors of senescence are predisposed to cancer. OIS derived from Ras overexpression, BRAF mutation, and/or PTEN deletion, acts through additional signaling pathways to activate p19^Arf^ and p16^Ink4A^ common effectors of senescence (*43*). Moreover, some progeroid syndromes with elevated senescence show a high incidence of tumors, with many SASP factors having pro-tumorigenic properties (*43*). Senescent cells during embryonic development were shown to re-enter the cell cycle after birth, proliferate, and contribute to epithelial cell fate (*44*), therefore, programmed cell senescence is likely to influence cell fate decision, including reciprocal senescence and apoptosis cell fate decisions, through balancing the consequences of INK4 and p53 mediated senescence and apoptosis decisions, among other senescence physiological roles (*6*). The fate of senescent cells may be context-dependent; yet several lines of evidence raise the possibility that senescence and apoptosis are simultaneously engaged in stress responses. How cells determine the outcome between senescence and apoptosis is less clear. It may be the nature and degree of stress and/or the levels of INK4 proteins that help decide cell fate. We observed a complementary tumor suppression between Tp53 and Ink4ab in fish. This observation raises the possibility that INK4-products constitutes a condition compound haploinsufficiency, where loss of just the right amount of INK4 tumor suppressors and p53 signaling are necessary for tumor formation, as suggested from biological (*45*) and mathematical data (*46*), offering a failsafe system designed to orchestrate a compound genetic redundancy continuum between senescence and apoptosis to ultimately maintain tumor suppressor functions.

Tp53-knockout mice lack phenotypic changes but are prone to tumor formation at 6 months of age. Similarly, a fraction of mice with knockout of Cdkn2a (encoding p16^Ink4A^ and p19^Arf^) develop sarcomas or lymphomas at 5–9 months of age (*3*). Both models, however, retain other INK4 family members, principally p15^Ink4B^, which fulfil a critical tumor suppressor (*36*) and senescence (*11*) functions. Moreover, both models recapitulate a non-tissue specific compound defect of senescence and apoptosis: mice with Tp53 knockout and, to a less extent, Cdkn2a knockout mice have impaired DNA repair and thus, accumulate DNA damage. Their phenotype complicates the assessment of the selective contribution of senescence to cell fate and tumor suppression in these models. Our studies revealed that deficiency of p19^Arf^ in mouse HSCs, when combined with p16^Ink4A^ and p15^Ink4b^ deficiency, not only limits tumor suppression, but also enhances senescence-like cell fate phenotype characterized by enriched tissue stress alarmin signal Hmgb2 and expression of immune checkpoint and SASP targets Cxcl13, Ccl5, Tigit, Gzmb, and lag3 in a hypersecretory state, accompanied with uncontrolled cell proliferation with increased Pcna and Mki67, and defective differentiation, leading to disturbed BCZ, scattering of T-cells, and activation of OIS in spatially resolved single splenocytes. These phenotypes suggest that cellular senescence, either by directly altering the immune system fitness or indirectly through the modification of the tumor environment (TME), may generate a tumor-permissive, chronic inflammatory microenvironment to shelter incipient tumor cells, therefore permitting immune-unchecked growth and proliferation (*30*) and favoring the accumulation of tumor cells that are more resistant and more prone to evade the immune system (*47*).

Since zebrafish presented with a single INK4 ortholog of human p15^Ink4b^/p16^Ink4a^ and p14^ARF^, this offered a model system with unique advantages to study cell cycle and senescence regulation. Similarly, our murine studies by modeling the compound loss of these three INK4 proteins in mouse HSCs revealed the senescence-associated pro-tumorigenic and hypersecretory phenotypes in mouse splenocytes. Senescent cells can arise throughout the lifespan and, if persistent, can have deleterious effects on cell differentiation and immune cell functions, due to their secretory profiles. Altering senescent cell fate (*48*) and/or targeting persistent senescent cells by enhancing immunoscenescence surveillance (*18*) and the use of small-molecule senolytic drugs (*49*) all would have promise, only when guided by an improved understanding of senescent cell effects on progenitor cell differentiation, immune cell function and regeneration, at the single cell level, for ultimately offering new strategies for cancer prevention and therapy.

## Methods summary

### Experimental design

The animal models and experimental conditions are fully described (*51*). All animal experiments were performed in accordance with the institutional animal care and use committee–approved protocols at Rutgers University and University of Colorado Anschutz (#01209). Monitoring of the phenotypes of zebrafish with ink4ab and/or tp53 deficiency, histological analyses, IHC, and radiation studies were based on established protocols (*33*) as described previously (*51*). Female p15^Ink4b^/p16^Ink4a^ mutant mice were rederived in-house and crossed Vav-Cre (*37*) hemizygous males (Jax #035670) to generate Ink4ab/Arf^+/-^Vav-Cre^+^ or Ink4ab/Arf^-/-^Vav-Cre^+^ as described (*51*).

### Identification and functional analyses of the evolutionary conserved inkab in Danio rerio

The ankyrin consensus sequences (*2*) were used identify the zebrafish ink4ab gene in the zebrafish genome. UniPort and phylogenic analyses were performed as described previously (*51*). Determination of ink4ab expression in embryonic and adult fish tissues were based on (*40*) as described in detail (*51*). The danio rerio zebrafish line with a retroviral insertion in the ink4ab upstream regulatory region was identified using retroviral tagging (*22*) for screening of a retroviral insertional mutagenesis library (*24*) using LTR sequences (fig. S6C and fig. S7). Carriers for the viral insertion mutation were used to establish the ink4ab insertional mutant line and compared to oligos that interfere with splicing of exon 2 of *ink4ab* (*23*). Cell cycle analyses and senescence detection in danio rerio were based on (*40*) and as described in detail (*51*).

### Identification of senescence cells in response to oxidative and oncogenic stimuli

MEFs were generated and maintained for senescence induction as described previously (*51*). Flow cytometry data were acquired using a five-laser BD Biosciences LSR and analyzed using FlowJo version 10.10 software. Detection of senescence-associated β-gal was based on optimized use of a fluorescent substrate (fig. S11, A to C) and small molecule (GLF16) (*28*), as fully described (*51*).

### Single cell spatial studies of splenocytes from conditional mouse models

CosMx Spatial Molecular Imager (SMI) (Nanostring) was used to detect thousands of RNA transcripts and proteins in formalin-fixed paraffin-embedded (FFPE) tissue sections at subcellular resolution (*39*). The procedure involves staining of the samples with a combination of RNA probes and oligonucleotide conjugated-antibodies that are detected in consecutive rounds of UV-photocleavable reporter hybridization (*39*). CosMx data were collected using the AtoMx software version 3.0.2 (Nanostring).

### Computational analysis

CosMx mouse universal cell characterization panel with the standard morphology panCK/CD45 protein markers were used in two slides with two technical replicates per tissue. High-resolution spatial capture (0.2 μm/pixel) across >6 field of views (FOVs) were processed for each tissue, filtered with a custom pipeline (*51*), and log library size normalized. Processed transcriptomic and morphology proteomic datasets were analyzed. Seurat (v5.1.0) R (v4.4.1) custom scripts were used for cell typing and visualization. Samples that clustered separately in initial Uniform Manifold Approximation and Projection (UMAP) visualizations, indicative of global shifts in gene expression, were integrated with Seurat CCAIntegration method to ensure comparability across cell types. Cell clustering was performed using Seurat’s shared nearest neighbor (SNN) modularity optimization-based clustering algorithm. The resolution parameter of 1.7 was adjusted to optimize the granularity of the resulting clusters. Cell types were assigned based on cluster-defining markers identified through FindMarkers, metacell inference, and literature references, and validated with the expression patterns (fig S18A). Cell-state annotations were confirmed with FindMarkers, metacell inference, and multimodal integration, using Seurat v5.0.1 package for cell typing, as previously described (*50*) and fully described (*51*).

Differentially expressed genes (DEG) analysis was conducted between WT, heterozygous, and homozygous mouse splenocytes using the FindAllMarkers function with a log-fold change minimum threshold of 0.25 and adjusted p-value < 0.05, among other analyses fully described in (*51*).

### Statistical analysis

Data are presented as means ± SD unless otherwise specified. Seurat FindMarkers with the default Wilcoxon rank-sum test was used for cell marker identification. The Benjamini-Hotchberg procedure was used to make corrections for multiple testing. A P value of <0.05 was used to denote statistical significance unless specifically indicated. Statistical analyses were performed using Prism version 10.0 (GraphPad).

## Acknowledgments

We thank Rutgers Cancer Institute of New Jersey Biorepository Services and Tissue Analytical Services (Shafiq Bhat, Lucyann Franciosa, Kelly Walton and Lei Cong) for assistance with tissue acquisition, histology, immunohistochemistry and sample processing. We thank Christine Archer for maintenance of zebrafish lines and Alison Hinojosa for assistance with mouse breeding.

## Funding

National Cancer Institute (5R01CA226746 to HES).

NCI-CCSG (P30CA072720 to SL).

NCI-CCSG (P30CA046934 to RS).

## Author contributions

Conceptualization: LL, SD, JK, TN, HES

Methodology: LL, SD, SY, AS, YZ, PV, KF, KJ, JK, CA, TN

Investigation: LL, SD, SY, AS, PV, KF, KJ, JK, CA, TN

Visualization: LL, SD, SY, AS, PV, KF, KJ, WW

Funding acquisition: HES

Project administration: SRP, HES

Supervision: JK, SL, BG, AVB, WW, SRP, TN, HES

Writing – original draft: HES

Writing – review & editing: LL, SD, SY, AS, YZ, PV, KF, KJ, JK, CA, SL, BG, AVB, WW, SRP, TN, HES

## Data and materials availability

All data are available in the main text or the supplementary materials.

Codes and materials used in the single cell spatial analysis will be available for the purposes of reproducing or extending the analysis at: https://github.com/orgs/SabaawyLab/repositories

