## Supplementary data for "Evolutionary conserved reciprocal senescence and tumor suppressor signals limit lifetime cancer"

**The PDF file includes:**

Materials and Methods
Figs. S1 to S21
Tables S1 to S8
References 51 to 65

### Materials and Methods

Zebrafish and mouse models

Zebrafish were raised and maintained under standard conditions and utilized according to the US Institutional Animal Care and Use Committee (IACUC) guidelines at Rutgers University and University of Colorado Anschutz (#01209). Monitoring of the phenotypes of zebrafish with ink4ab and/or tp53 deficiency, histological analyses, IHC, and radiation studies were based on established protocols (*1*). Female $p15^{Ink4b}/p16^{Ink4a}$ mutant mice were rederived in-house and crossed with Vav-Cre (*2*) hemizygous males (Jax #035670) to generate $Ink4ab/Arf^{+/-}Vav\text{-}Cre^{+}$ or $Ink4ab/Arf^{-/-}Vav\text{-}Cre^{+}$. More than 6 generations of mice were utilized in our studies with consistent phenotypes.

Gene prediction tools

Computational screening of the ink4ab gene was conducted using the zebrafish genome database (www.sanger.ac.uk/cgi-bin/blast/submitblast/d.rerio). The putative zebrafish *ink4ab* gene was retrieved with a BLASTP search using ankyrin domain amino acid sequences shared among the human INK4 family (Ensembl accession number ENSDARP00000054208). Genomicus, a database and webserver for comparative genomics in eukaryotes was used to graphically display the synteny conservation of the INK4 loci between multiple genomes (*3*). A total of 1029 extant genomes and 621 ancestral reconstructions from all eukaryotes kingdoms available in Ensembl and Ensembl Genomes databases were used. The Protein Sequence Alignment tool was used in MacVector to compare sequence and structural homology between Danio rerio Ink4ab, and other INK4A and INK4B orthologs from other species. MacVector software was also used to create a neighbor-joining phylogenetic tree. PHYML was used for estimating the maximum likelihood phylogenies from DNA and protein sequences (*4*). With PHYML, we used nonparametric bootstrap and estimation of various evolutionary parameters, in order to perform a comprehensive phylogenetic analysis on large datasets at http://atgc.lirmm.fr/phyml.

The ankyrin consensus sequences which are required for the interactions between $p15^{Ink4b}/p16^{Ink4a}$ and CDK4/6 proteins have been described (*5*), these consensus sequences were used to confirm evolutionarily constrained regions (ECRs) for ankyrin domains. ECRs that harbor ankyrin repeats are a hallmark for sites of critical importance for an INK4 protein's structure or function (*6*). The ankyrin consensus ECRs were inferred by comparing the amino acid sequences from multiple protein homologs in the context of the INK4 evolutionary relationships using Aminode (*6*). Aminode was pre-loaded with the results of the analysis of the whole human proteome compared with proteomes from 62 additional vertebrate species. Profiles of the relative rates of amino acid substitution and ECR maps of INK4 proteins were used mapping ARF related sequences at http://www.aminode.org.

ClustalX features were then utilized to generate a graphical interpretation of the alignments of the zebrafish ink4ab polypeptide sequence with human, mouse, rat, fugu, and Western clawed frog (xenopus) INK4A and INK4B orthologs while mapping the ankyrin domains using ClustalX features (*7*).

PCR cloning and sequencing of the *ink4ab* gene

The synthesized cDNA was used as a template for PCR to clone *ink4ab* gene from two cDNA libraries generated from the 24 hours post-fertilization (hpf) zebrafish embryos and brain tissues of adult male and female zebrafish. The primers used for each member of *ink4ab* were designed to target the full-length of *ink4ab*. The PCR products were cloned into pGEM-T easy vector (Promega) or PCS2p$^+$ (Addgene) for *in vitro* transcription and sequencing. The cDNA clones were sequenced by GeneWiz, Inc.

RNA isolation from zebrafish embryos

Embryos were dechorionated, deyolked and washed with embryo media. Embryos were centrifuged and the supernatant was removed. The pelleted embryos were then frozen on dry ice and homogenized using a pestle in 250μl of Trizol Reagent (Invitrogen). Pestles were rinsed with an additional 250μl of Trizol and 100μl of Chloroform was added to each tube. Tubes were shaken vigorously by hand for 15 seconds and then incubated at room temperature for 2-3 minutes followed by centrifugation at 12,000 g for 15 minutes at 4°C. The top aqueous layer containing RNA was carefully collected and transferred clean Eppendorf tubes. RNA was precipitated by adding 250μl of isopropanol, incubating samples at room temperature for 10 minutes or at 4°C for 30 minutes, followed by centrifugation at 7,500 g for 5 minutes at 4°C. RNA pellets were washed with 75% ethanol, re-centrifuged, and then air-dried pellets were dissolved in RNase-free water.

RT-PCR

Total RNA was isolated from embryos using Trizol reagent, and 1 μg of RNA was used for reverse transcriptase reaction with RevertAid (Promega). PCR was performed using GoTaq Polymerase (Promega) with primers targeting *beta-actin* or zebrafish *ink4ab* cDNA.

Antisense RNA Probe preparation

RNA was isolated from zebrafish embryos using Trizol and reverse transcribed with Fermentas RevertAid cDNA Synthesis. To determine the expression pattern of *ink4ab* during development, the ink4ab cDNA was synthesized from mRNA isolated from 24 hpf embryos, amplified by PCR. The ink4ab cDNA was subsequently cloned into a TOPO vector to introduce flanking restriction enzyme sites. The ink4ab cDNA was cloned into pGem-T-Easy vector (Promega) and linearized at a site before a T7 promoter sequence for *in vitro* transcription. The *in vitro* transcribed ink4ab antisense mRNA was labeled with digoxigenin (DIG) to detect *ink4ab* expression by *in situ* hybridization. The cDNA was then reverse transcribed and labeled with digoxigenin (Roche) to prepare an antisense RNA probe. The labeled probe was subjected to formaldehyde agarose gel electrophoresis to determine concentration and quality of RNA.

*In situ* Hybridization

To prepare embryos for *in situ* hybridization, embryos were fixed in freshly made 4% paraformaldehyde for 4 hours at room temperature or overnight at 4°C. Embryos were subsequently washed in PBS three times for 5 minutes each and then dehydrated in 100% methanol at -20°C for at least an hour. Embryos were then rehydrated in graded methanol/PBST (PBS plus 0.1% Tween 20) and washed three times in PBST. Embryos were then digested with proteinase K (10μg/ml in PBST) at room temperature for 30 minutes or less depending on the age, washed in PBST and fixed in 4% paraformaldehyde for 20 minutes. Embryos were then prehybridized at 68°C for 3 hours followed by incubation in the probe overnight with gentle rocking. The next day, embryos were washed and blocked with 5% lamb serum for 2 hours and incubated in digoxigenin antibody overnight at 4°C. Embryos were washed in PBST several times followed by staining with Nitro Blue Tetrazolium and 5-Bromo 4-Chloro3 Indolyl Phosphate. The reaction was stopped by removing the staining solution and washing the embryos in 3 quick washes of PBST followed by a low pH (3.0) PBS wash for 5 minutes. Embryos were fixed again in 4% PFA for 30 minutes and washed 3 times with PBST.

Generation of ink4ab mutant zebrafish

Zebrafish lines from the retroviral insertional mutagenesis screen (*8*) were examined for retroviral insertions into the genomic region of the *ink4ab* gene. Zebrafish were genotyped for verification of the mutation using two PCR assays. The first PCR included primers specific for viral long terminal repeat sequences (LTRs) to determine that a viral insertion occurred. The second PCR was performed using an aliquot of the first PCR product and primers specific for the zebrafish genomic DNA to amplify the *ink4ab* promoter sequence flanked by the viral LTRs (fig. S6C and fig. S7A). Five female fish and four males were identified as carriers for the viral insertion mutation and were used in heterozygous intercrosses to generate homozygous ink4ab mutants (fig. S7B). Morpholino oligomers (MO) are synthetic oligonucleotides composed of chains of about 25 subunits that are similar to DNA and RNA oligonucleotides, except that they have a morpholine ring rather than a ribose ring making them more stable, less toxic and resistant to nucleases. MOs were used to induce ink4ab gene knockdown by blocking translation initiation (*9*) or by interfering with splicing (*10*). Multiple MOs with distinct design features were first screened to identify antisense MOs to transiently knock down (KD) *ink4ab* by using complete pre-mRNA sequence for defining suitable targets, validating the targeting of *ink4ab* by RT-PCR analysis of splice modification in a beta-actin splice model system, and simultaneously blocking the U1 and U2 small nuclear Ribonucleoprotein (snRNP) binding sites and splice-junctions (*11*). It was next confirmed that embryos injected with control or target MOs do not show a systemic immune response or off-target splicing defects as previously demonstrated in xenopus (*12*). The MO dosage was optimized, and incubation temperatures were increased to achieve consistent dose-dependent *ink4ab* KD (fig. S6D) and mimic the effects of the ink4ab insertional mutant zebrafish line.

Fin clipping and DNA extraction

Adult zebrafish were anesthetized using tricaine for no more than 5 minutes. Fish were placed on a clean surface and excess water was removed using kimwipes. The fins were spread so that they were flat and then cut by gently rolling of a scalpel across the fin. Fins were then placed in 100µl of DNA extraction buffer consisting of 400mM Tris-HCl, 5mM EDTA, 150mM NaCl, 0.1% Tween-20 and 1mg/ml proteinase K. Samples were incubated overnight at 60°C with shaking at 300 rpm. An amount of 200µl of water was added to each sample and then samples were heated to 95°C for 10 minutes to inactivate the proteinase K and then centrifuged for one minute at 17,000 g. Following that, 2µl of the supernatant were used for PCR genotyping.

Microinjection of plasmids, mRNA and morpholinos

One-cell-stage zebrafish embryos were injected with 2.5nl containing 1-3.3ng cdkn2a/b mRNA and/or MO diluted in 10% Phenol Red. MOs targeting the *ink4ab* ATG region

(5'CAGTTCATCCTCGACGTTCATCATC-3') or the 3' splice site of exon 1 (5'-AAAGCGCGTCTAAACCTACCTGTAT-3') were synthesized by Gene Tools.

A p53 MO (5'-GCGCCATTGCTTTGCAAGAATTG-3') was used at 4.5 ng/embryo.

Quantitative RT-PCR

Total RNA was isolated from embryos after homogenization, using Trizol Reagent (Invitrogen). cDNA was reversed transcribed using Oligo(dT)18 primers and M-MuLV reverse transcriptase (Fermentas) according to the manufacturer's instructions. Gene expression was determined in a Q-RT-PCR reaction using a custom TaqMan Gene Expression Assay (Applied Biosystems), in the automated Applied Biosystems StepOne Plus RT-PCR Real-Time Thermal Cycler. Gene expression was measured in triplicates across a set of three biological replicates. Beta-actin was used as an endogenous control and reference gene for sample normalization. Relative quantification and calculation of the range of confidence were performed using the comparative ΔΔCT method, as previously described (*13*).

Cell Cycle Analysis

The design of cell cycle analyses experiments in zebrafish embryos was completed as previously described (*14*). Embryos were dissociated at 48 hpf using reagents and steps that were previously described (*15*). Cells were fixed in 80% ethanol and washed in PBS. Cells were stained with 2µg/ml propidium iodide and subjected to flow cytometry. FlowJo software was used to analyze the cell cycle profiles.

Senescence induction and senescence associated β-galactosidase (SA β-Gal) Assay

Embryos were placed in either egg water alone or egg water containing 150µM hydrogen peroxide at 6hpf to induce oxidative stress ($O_2S$). Embryos were fixed in 4% paraformaldehyde at 6 day post-fertilization (dpf), washed in PBS at pH 7.4, followed by PBS pH 6.0 for one hour each. Fixed embryos were then stained with 2mg/ml x-gal (Sigma) overnight at 37°C and washed with PBS at pH 6.0. Embryos were imaged and staining was quantitated using Adobe Photoshop and ImageJ as previously described (*16*) and recently validated *(17),* with a few modifications. For image processing and to remove any potential background staining, the embryo proper was cropped using the selection tool in Adobe Photoshop. Cropped images were saved as TIF files and opened in ImageJ. The β-gal staining intensity was measured using the average intensity density of hue saturation.

Two-step PCR cloning of pMam-EGFPink4ab

An optimized PCR for amplification of ink4ab cDNA was performed. KOD Xtreme was used with cDNA from adult brain tissue as template for amplification of ink4ab cDNA by PCR. ink4ab cDNA was digested with BamHI and EcoRI. pEGFPc1 was digested with BglII and EcoRI. After ligation and transformation, a positive clone was confirmed using AfeI and EcoRI digestion and KasI digestion. Plasmids pEGFPc1-EGFPink4ab were digested with NheI and SalI then the EGFPink4ab 1.15kB fragment was gel purified. EGFP-ink4ab was inserted into pMam-Neo between NheI and SalI sites. Digestion of two clones with EcoRI and NcoI was used for further confirmation of the correct clones, followed by sequencing of the insert region to further confirm correct sequences and orientation.

Preparations of embryo lysates for Western blotting

Embryos were dechorionated, deyolked, and washed in embryo media three times. All media was removed. Embryos were frozen on dry ice and then homogenized using a pestle. Lysis buffer consisting of RIPA buffer supplemented with protease inhibitors (Mammalian Protease Arrest) was added at a concentration of 2-10μl of lysis buffer per homogenized embryo and then incubated on ice for 30 minutes. Lysates were sonicated at 40Hertz for three 10-second intervals and cooled on ice in between intervals. Lysates were incubated on ice for an additional 15 minutes and then centrifuged at 12,000 g for 5 minutes. Laemmli buffer supplemented with β-mercaptoethanol (BME) was added and then lysates were boiled at 95°C for 5 minutes. Samples were placed on ice and then subjected to protein agarose gel electrophoresis.

Western blotting

Lysates from NIH3T3 cells were electroporated with pMam-Neo, pMam-mp15, or pMam-EGFP-ink4ab and treated with 1μM dexamethasone or 0.04% ethanol (solvent). The protein concentrations were determined using a BCA assay. In each lane, 40μg of protein was subjected to gel electrophoresis on a 10% Tris-glycine-SDS gel. Proteins were transferred for one hour at 0.35 constant Amps onto a 0.45μm PVDF membrane by a wet-transfer system in transfer buffer containing 20% methanol. Membranes were processed with methanol, dried, and then re-wet with methanol prior to blocking in 5% milk for one hour. The membranes were incubated overnight at 4°C in primary antibody. The membranes were then washed and anti-rabbit-HRP or anti-mouse-HRP were used as secondary antibody at a concentration of 1:10,000 for one hour at room temperature. The membranes were stripped using the low pH method (glycine-HCl SDS), blocked in either 5% milk or BSA for one hour, followed by a one-hour incubation in primary antibody, washed, one hour incubation in secondary antibody, and a final wash. Anti-GFP (Av-GFP JL-8, Living Colors) was used to detect zebrafish ink4ab-GFP fusion protein expression because there is a paucity of validated antibody reagents for use against zebrafish proteins (*18*) and specifically no zebrafish ink4ab antibody currently available and human or murine antibodies against $p15^{Ink4b}$/$p16^{Ink4a}$ lack cross reactivity. The predicted size for the EGFP-ink4ab fusion protein is approximately 45kD.

Cell culture and examination of KRas induced senescence in murine cells

NIH3T3 mouse embryonic fibroblasts were maintained in Dulbecco's Modified Eagles Medium (DMEM) supplemented with 10% Colorado calf serum and 1% penicillin/streptomycin. Cells were passaged every 5 days at subconfluency to maintain the contact inhibition property of NIH3T3 cells. 293T human embryonic kidney cells were maintained in DMEM supplemented with 10% fetal bovine serum plus 1% penicillin streptomycin and passaged every 4 days. Murine cells were transfected with the following plasmids using electroporation or Lipofectamine LTX (Invitrogen) (Table S8). Cells transfected with pBabe plasmids (pBabe-puro and pBabe-$HRas^{V12}$) were selected with puromycin. The pMam plasmid transfected cells were selected with Geneticin (G418). Puromycin selection was optimized to an optimum selection concentration starting at 400μg/ml G418, while 800μg/ml resulted in better selection (100% of non-transfected cells died and colonies remained for transfected cells after one week). The inducible expression of either mouse $p15^{Ink4B}$ or zebrafish ink4ab was used to prevent the continuous cellular senescence of cells, when they are stably transfected with either mouse p15 or zebrafish ink4ab. After two weeks of G418 treatment, positive clones were selected and plated into 12 well plates for growth analysis. Cells transfected with either pMam-Neo (vector), pMam-mp15 (mouse p15), or pMam-

EGFP-ink4ab (zebrafish ink4ab) alone or in combination with pBabe-$HRas^{V12}$ (oncogenic HRas), treated with the steroid dexamethasone to induce expression of mp15 and ink4ab or with ethanol as a control. Cells were collected and counted manually using a hemocytometer every other day for a week.

Next, the cell cycle of 293T cells transfected with either pEGFP-c1 (vector control) or pEGFP-ink4ab alone and in combination with pBabe-$HRas^{V12}$was analyzed. At 48 hours after transfections, cells were sorted to enrich for GFP positive cells. The sorted cells were incubated at 37ºC for a day to allow recovery from cell sorting. The following day, cells were fixed and stained with propidium iodide, and the cell cycle was analyzed using flow cytometry. Similar to results of pRB phosphorylation changes under basal conditions, differences in the cell cycle profiles between cells transfected with pEGFPc1 or pEGFP-ink4ab alone were examined. Upon oncogenic stimulation, there was a modest increase in the percentage of cells in G1 and a decrease in the percentage of cells in S-phase of the cell cycle observed in cells expressing both ink4ab and oncogenic HRas.

Mammalian cells cease proliferation in culture when spatially confined by their neighbors, a phenomenon called contact inhibition. During contact inhibition, cells do not undergo senescence (*19*). Moreover, immortalized mouse embryonic fibroblasts (MEFs) lose their strong contact inhibition over time. Therefore, in addition to assessing senescence in the traditional NIH3T3 clones, a new clone (3T3.2, ATCC) derived by single cell cloning and having the validated feature of restored contact inhibition was utilized to study senescence in response to oncogenic Ras activation. We confirmed $Kras^{V12}$ activation in either 3T3.2 cells or $p16^{Ink4A-/-}$/$BMI1^{+/-}$ MEFs resulted in increased senescence-associated β-gal, confirmed by detecting an optimized fluorescent substrate (fig. S11, A to C) and the results were validated with a second senescence assay using a fluorescent small molecule (GLF16), which was recently developed and shown in elegant studies to detect senescent associated β-gal positive cells as previously described (*20*).

<u>Campothecin treatment and detection of apoptotic cells in zebrafish embryos</u>

Embryos were treated with campothecin (CPT). Campothecin is a topoisomerase inhibitor, which induces single-stranded DNA breaks that ultimately lead to double-stranded breaks due to collapse of replication forks. As a result of the increased double-stranded breaks, apoptosis is activated. Embryos were treated with 50nM, 200nM, or 500nM CPT and ink4ab expression was measured using RT-PCR. It was first determined that *ink4ab* mRNA expression increases in WT embryos in response to a 6-hour CPT treatment (fig. S10, B to C). Acridine orange staining was then used as a method for whole-mount detection of apoptotic cells in live embryos. To determine if the apoptosis activity in *ink4ab* mutant embryos is mediated through the p53 pathway, embryos were injected with a p53 MO and measured apoptotic cells in embryo tails. Knockdown of p53 with MOs dramatically reduced the number of apoptotic cells in both untreated and CPT treated *ink4ab* mutant embryos. This suggests that *ink4ab* mutants have increased activation of p53-dependent apoptosis. Embryos were dechorionated and incubated in 50nM CPT for 6 hours and 28.5°C. Embryos were rinsed quickly 3 times with egg water and then incubated in 2µg/ml Acridine Orange. After 30 minutes, embryos were washed with egg water 3 times for 5 minutes each and then imaged on a fluorescent microscope.

Histologic Evaluation of Solid Tumors

In order to establish a correlation between ink4ab deficiency and the presence of a neoplasm, fish were examined histologically. Adult fish were fixed in Dietrich's fixative with gentle rocking for a minimum of five days. Fish were washed in PBS and then in 70% ethanol after fixation. The entire fish were bisected longitudinally along the midline and both halves were processed for histologic evaluation to determine the extent of tumor invasion. Paraffin-embedded tissue sections (4μm) were used for H&E staining and immunohistochemistry. Antibodies used for immunohistochemistry include anti-PCNA (1:4,000, Sigma) and anti-phospho-histone H3 (PH3) (Ser10) (1:300, Millipore).

Gamma irradiation and assessment of melanocytes

A cohort of thirty WT and *ink4ab*$^{-/-}$ fish were subjected to 23Gy of gamma radiation as previously described (*1*) and monitored daily for the presence of tumors or depletion of health. Schmorl staining was performed on sections from adult fish with apparent melanocytic neoplasia as described (*21*). Paraffin sections were deparaffinized in xylene and rehydrated in descending concentrations of ethanol. The sections were incubated for 10 minutes with a working solution comprising 0.75% ferric chloride and freshly prepared 0.1% potassium ferricyanide in distilled water and washed in distilled water. Cells containing melanin are stained in blue. Nuclear Fast Red was used as a counterstain (2 minutes) before sections were dehydrated in ethanol.

Analysis of hematopoietic cells

Cytospins were generated using dissected spleen and kidney marrow cells from four one-year old wild type and *ink4ab* mutant zebrafish then stained with Giemsa. Spleens and kidneys were dissected from five WT and mutant fish, homogenized by passing cells through a 40 μM cell strainer, and washed in PBS containing 10% FBS. The cells were subsequently washed twice in PBS, stained with 7AAD for exclusion of dead cells, and subjected to flow cytometry using forward and side scatter to detect subpopulations of hematopoietic cells as previously described (*1*). FlowJo was used to analyze flow cytometry data.

Single cell spatial studies

The CosMx Spatial Molecular Imager (SMI)(*22*). To characterize the single-cell spatial distribution of splenic cell phenotypes, spatially resolved single cell transcriptome was analyzed in splenic sections from WT, in the Ink4ab/Arf$^{HSC+/-}$ and Ink4ab/Arf$^{HSC-/-}$ mouse spleen sections. The composition of the 1000-probe panel included 243 genes for cell typing, 269 genes for cell function, 435 genes for cell-cell interaction, and 46 senescence-associated targets (*23-25*). For CosMx SMI data, the mean number of transcripts per cell was 294 for control slides and 268 for Ink4 slides, with an average of over 100 transcripts per cell (as a threshold criteria) among 40 FOV. The probe detection sensitivity of the CosMx SMI was determined to be 1 to 2 copies per cell. Single-cell expression profiles were derived by counting the transcripts of each gene that fell within the area assigned to a cell according to the cell segmentation algorithm. An average of 4% of cells contained fewer than 20 total transcripts, which were omitted from further analysis, because this class of cells behaves anomalously when projected into lower dimensions (for example, uniform manifold approximation and projection (UMAP)). The readout process of the CosMx SMI technology is shown after decoding the bound probe barcodes, morphological, spatial, and cell-type specific information. Spatial features and mRNA cellular localization within the tissue section were collected using the CosMx software AtoMx (*22*). CosMx mouse universal cell characterization panel with the standard morphology panCK/CD45 protein markers were used in two slides with two technical replicates per tissue. High-resolution spatial capture (0.2 µm/pixel) across >6 field of views (FOVs) were processed for each tissue, filtered with a custom pipeline, and log library size normalized. The AtoMx workflow allows the semi-supervised clustering to annotate all cells with either a known cell type or a novel cell type. Cells grouped into novel cell types had consistent expression profiles within one cell type but did not match the expression profile of any known cell type in the initial reference profile. Because the novelty of the CosMx assay and lack of reference murine cell typing datasets, we attempted to validate the cell clustering through AtoMx by cross-matching human splenocyte lineage markers. However, since there are recently documented structural differences between the murine and human splenic immune system (*26*), most notably, the distinct organization of T-cell zone (TCZ) and B-cell zone (BCZ) follicles (*26*), and the dynamic and overlapping marker expression by different T-cell and dendritic cell (DC) subsets, it has historically been challenging to segregate T-cells and DCs into clear lineages (*26*), with the most widely used classification systems of splenic murine DCs for example as $CD4^+$, $CD8^+$, or double-negative ($CD4^-CD8^-$) DCs (*26*). To address these limitations, processed transcriptomic and morphology proteomic datasets were analyzed with a custom pipeline. Seurat (v5.1.0) R (v4.4.1) custom scripts were used for cell typing and visualization. Samples that clustered separately in initial UMAP visualizations, indicative of global shifts in gene expression, were integrated with Seurat CCAIntegration method to ensure comparability across cell types. Cell clustering was performed using Seurat's shared nearest neighbor (SNN) modularity optimization-based clustering algorithm. The resolution parameter of 1.7 was adjusted to optimize the granularity of the resulting clusters. Cell types were assigned based on cluster-defining markers identified through FindMarkers, metacell inference, and literature references, and validated with the expression patterns (fig S18A). Cell-state annotations were confirmed with FindMarkers, metacell inference, and multimodal integration, using Seurat v5.0.1 package for cell typing, following the general principles of single cell spatial cell typing, as recently described (*27*). Characteristic marker gene expression profile of each of the 14 cell type were utilized to assign cell identities to each cluster. Specifically, two clusters were undefined as others 1-2, and the following 12 cell types were identified: (1) B-cells (expressing Ighd and CD19); (2) plasma cells (expressing J-chain, Ighg1 and Xbp1); (3) T-cells (expressing Cd3d, Nkg7, Cd3e, and Cd8a); (4) neutrophils (expressing Lcn2, Ltf, and Hmgb2); (5) DCs (expressing S100a4 and Itgax); (6) monocytes (expressing C1qb, C1qa, C1qc, and Selenop); (7) macrophages (expressing Thbs1 and Myl9); (8) erythroid progenitors (expressing Hmgb2, Ube2c, Mt1, and Top2); (9) fibroblasts (expressing Acta2, Clu, Igfbp3, and Tagln); (10) granulocytes (expressing Mpo, Elane, and Prtn3); (11) plasmacytoid DCs (expressing Xbp1 and C1qb); and (12) plasmablasts (expressing Selenop and Clu). To analyze the genes expressed in the BCZ and TCZ clusters, the levels of differential gene expression (DGE) of senescence associated marker genes for annotation was conducted between WT, heterozygous, and homozygous mouse splenocytes using the FindAllMarkers function with a log-fold change minimum threshold of 0.25 and adjusted p-value < 0.05. After identifying that neutrophils, monocytes and erythroid progenitors are enriched in the  in our dataset, the DEGs in the spleens of Ink4ab/Arf$^{HSC+/+}$ and Ink4ab/Arf$^{HSC-/-}$ mice were identified. DEGs were selected based on two criteria: (| log2 (fold change) | > 1) and significance level (p< 0.05).

Statistical analyses

Statistical analyses of survival and comparative rates of tumor formation were performed by a qualified Biostatistician. For pair-wise comparison between groups, the Sidak multiple-comparison adjustment was used to control overall significance level at 5% and a Cox proportional hazard model was employed to determine statistical significance. For statistical analysis of all other data, standard deviations were calculated for averages from experimental replicates and standard error of the means were derived. Student T-tests were used, and significance was set with p-values of less than 0.05.

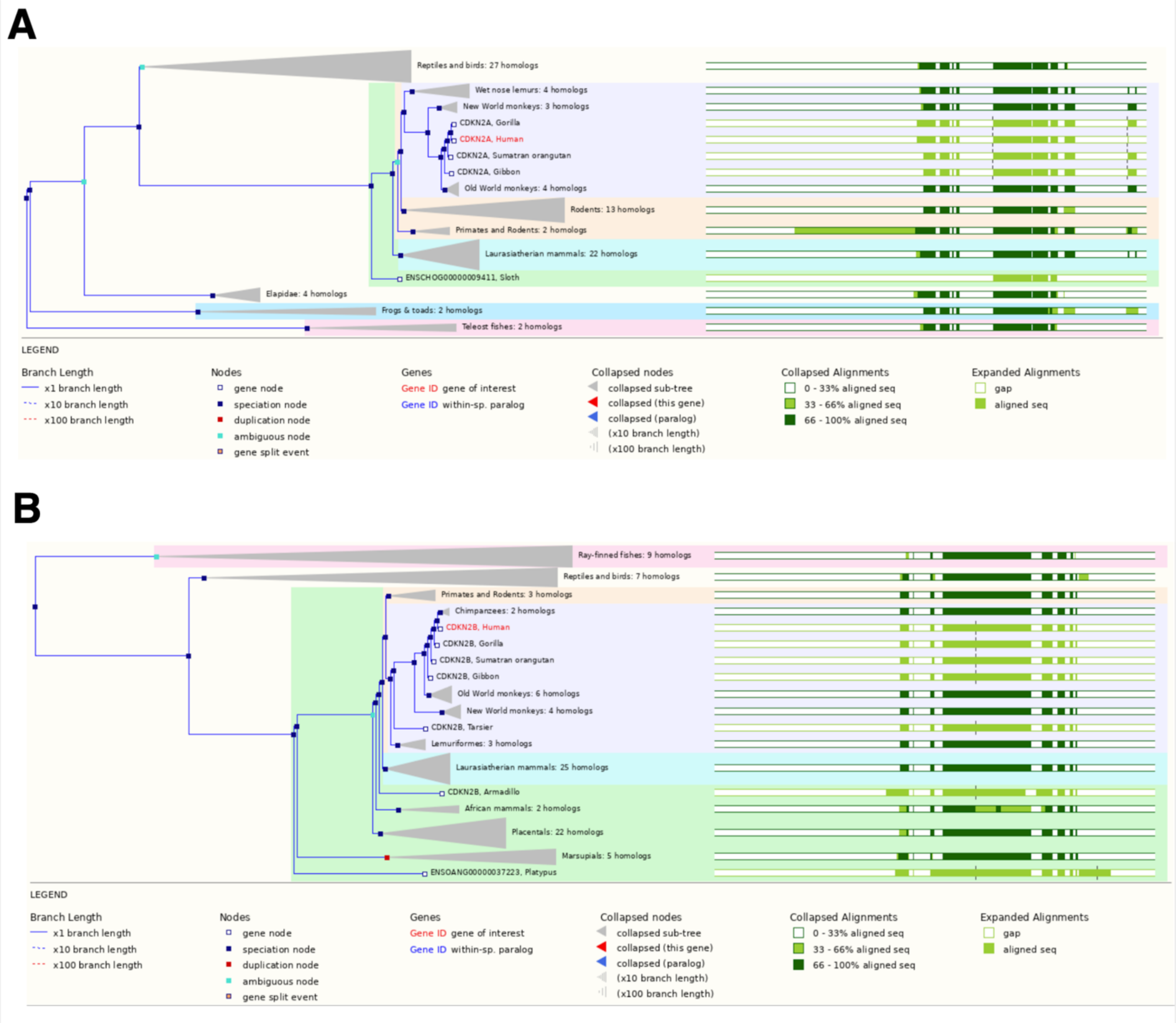


**Fig. S1. Phylogenetic analyses of INK4 genes.** (**A**) Phylogenetic tree of CDKN2A. (**B**) Phylogenetic tree of CDKN2B. Human genes are encoded by a red color. Gene ID are listed based on Ensembl Human dataset (GRCh37). Human p14$^{ARF}$ is an alternative transcript to CDKN2A.

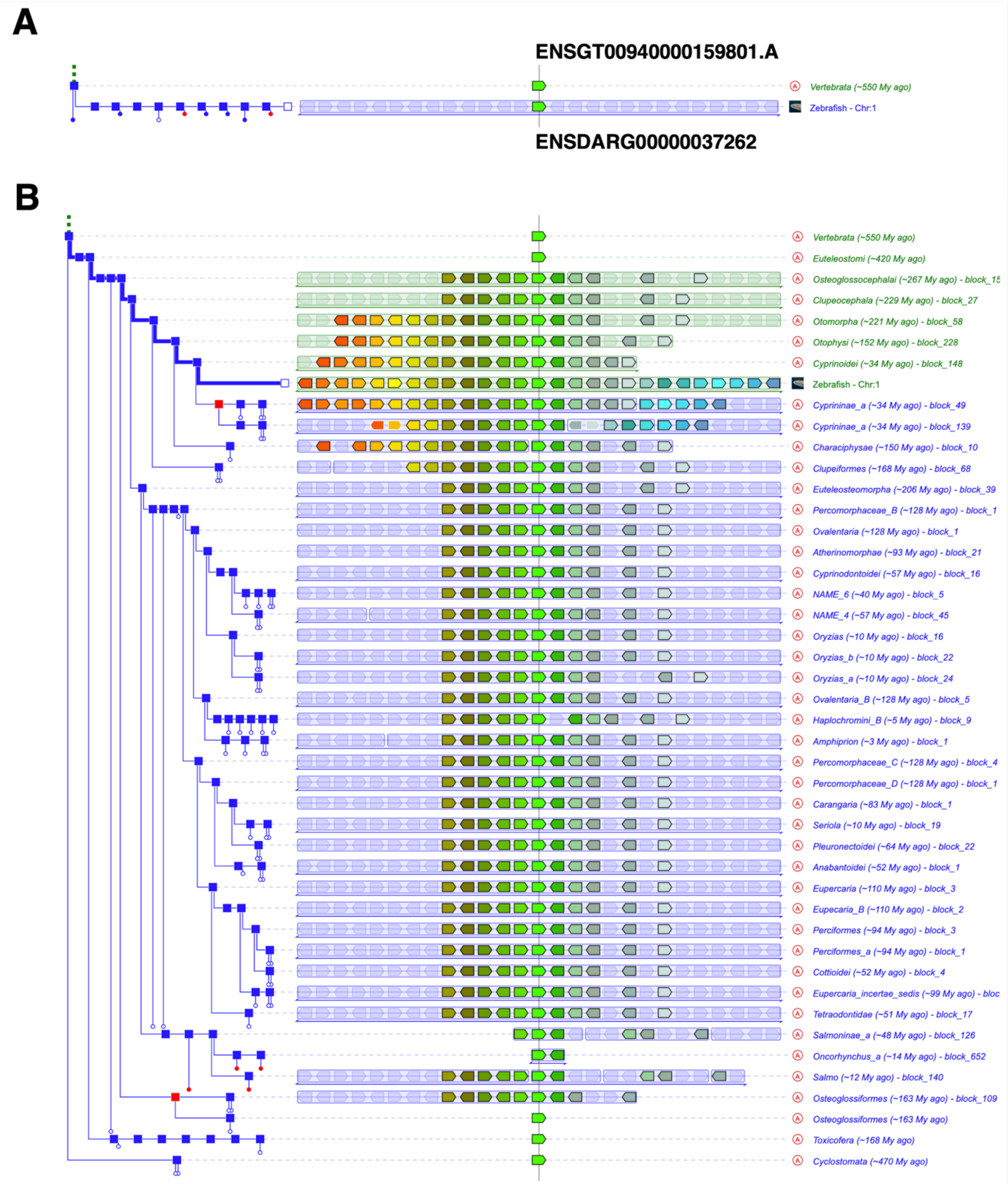


**Fig. S2. Phylogenetic origin and evolutionary sequence similarities of the zebrafish *ink4ab* gene.** (**A**) Compact AlignView of Genomicus showing an alignment between the zebrafish *ink4ab* gene contained within the genomic region of the reference gene ENSDARG00000037262 (bright green) and ancestral root Vertebrata ENSGT00940000159801.A. (**B**) PhyloView showing the order and orientation of respective orthologs in different genomes that share the same ancestral root with the zebrafish *ink4ab* gene at ENSDARG00000037262, the order of reference of neighboring genes, and the order of their respective orthologs and paralogs in different species that share the same ancestral "root" species. The tree on the left of the display is the phylogenetic tree (computed by Ensembl) of the zebrafish *ink4ab* gene shown in the middle that intersects the vertical line. Red square nodes represent duplication events of an ancestral version of the *ink4ab* gene used as query. Therefore, the Osteoglossiforms (~163 million years ago) and Cyprininae species (~34 million years ago) appear twice due to a duplication node shown as square (as in the second node along the red path). Orthologs of these genes in other species are shown in matching colors. Note that the neighboring gene to *ink4ab* shown in dark green is the *mtap* gene. Blue square nodes represent ancestral species leading from the same "root" ancestral species to orthologs and/or paralogs of the *ink4ab* gene used as query. Red square nodes represent duplication events of an ancestral version of the *ink4ab* gene used as query. Data generated using Genomicus v110-01 (*3*).

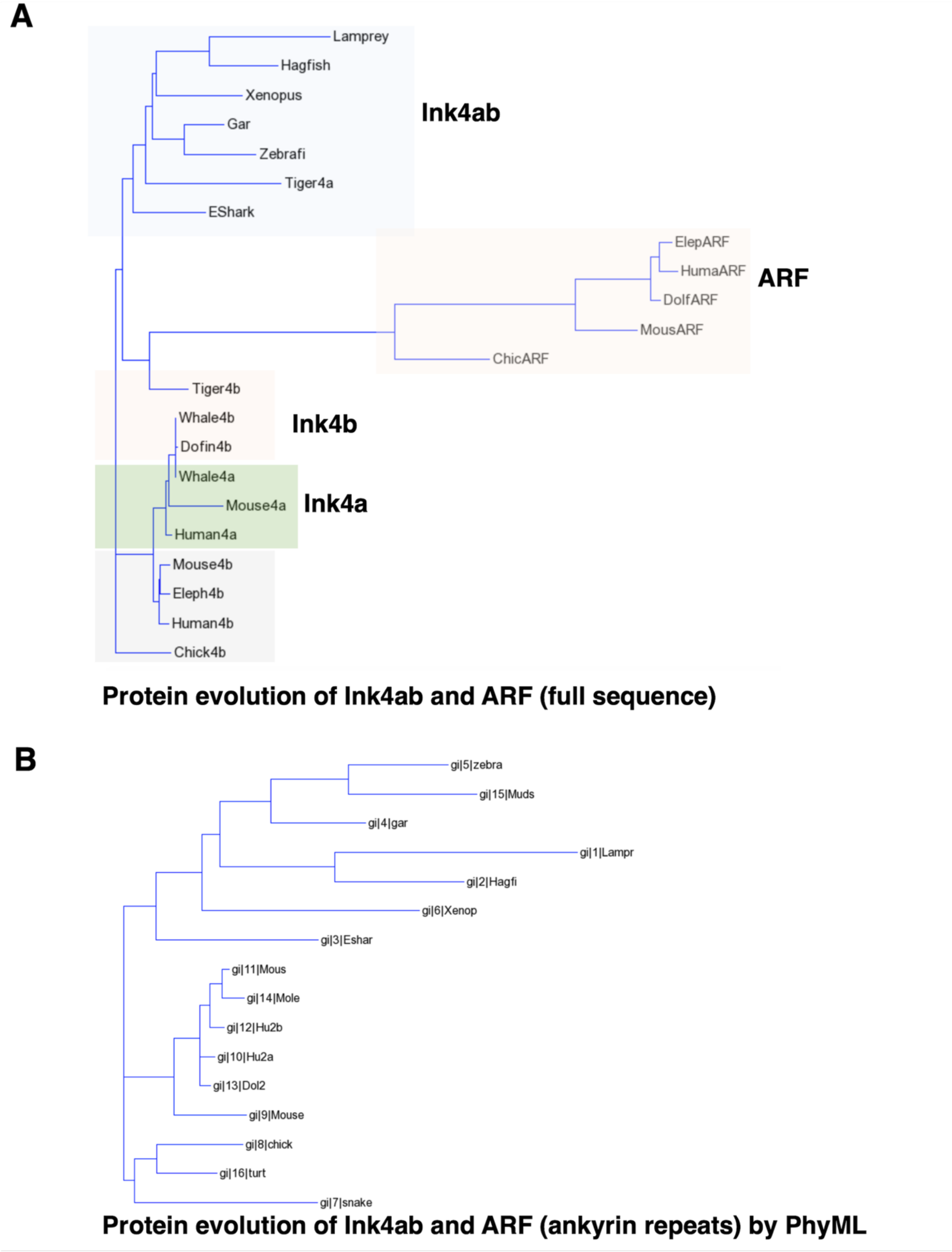


**Fig. S3. Protein evolution of INK4 proteins.** (**A**) Phylogenetic tree of Ink4a, Ink4b, and combined Ink4ab or ARF predicted amino acid sequences. (**B**) Phylogenetic tree of the conserved ankyrin domains generated using PhyML (*4*). Gar, alligator gar; zebrafi, danio rerio;

tiger, tiger snake; eshark, elephant shark; hagfi, hagfisf; muds, mudskipper; lamp, lamprey; xenop, western clawed frog; hum, human; mous, mouse; dol, dolphin; chivk, chicken; turt, desert turtle; snake, tiger snake.

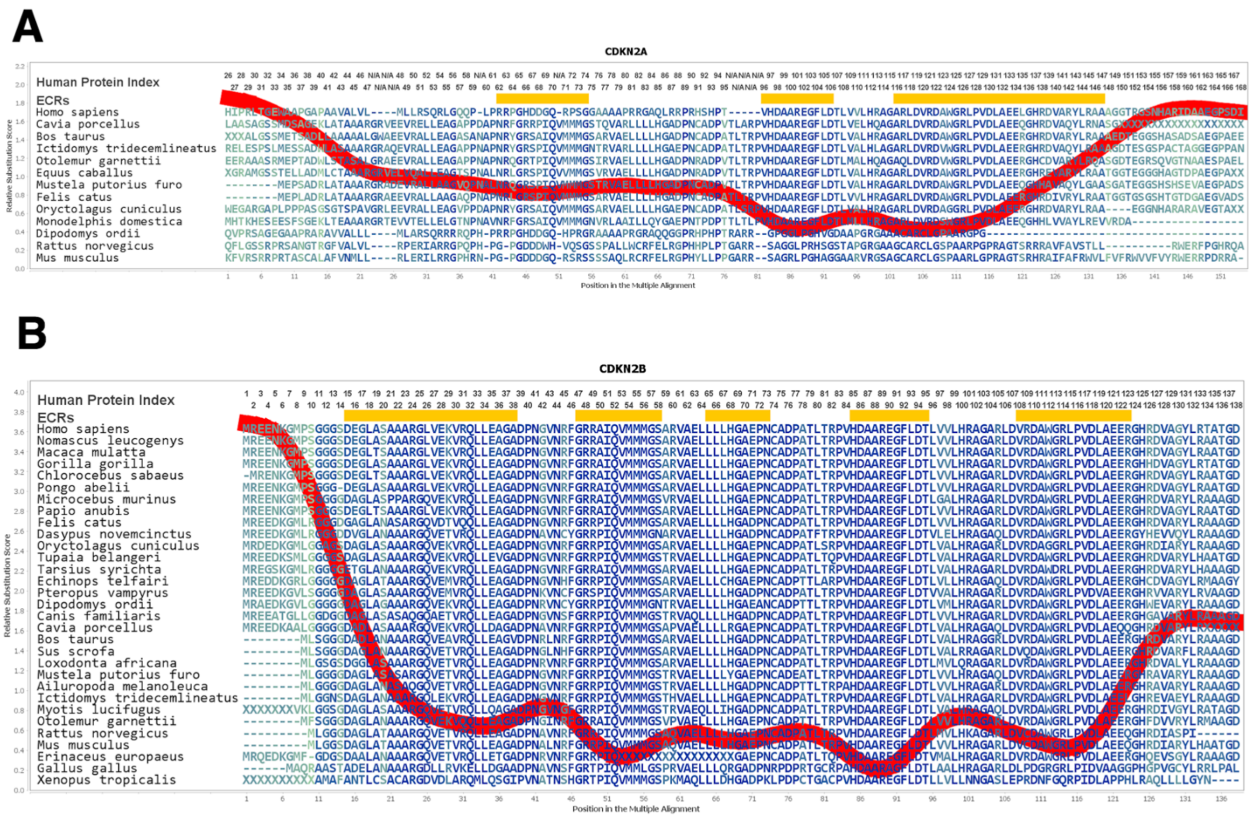


**Fig. S4. Evolutionary conservation of human CDKN2A (p16$^{INK4A}$) and CDKN2B (p15$^{INK4B}$) amino acid sequences and ankyrin domains.** (**A**) Multiple alignment and positions of amino acid sequences of CDKN2A (p16$^{INK4A}$) orthologs listed based on highest relative substitution scores compared to human sequences. (B) Multiple alignment of human CDKN2B (p15$^{INK4B}$). Evolutionary constrained regions (ECRs) are inferred by comparing the amino acid sequences from multiple protein homologs in the context of the evolutionary relationships that link CDKN2 proteins and are displayed with red line. Yellow lines indicate ankyrin domain (AD) in highly conserved regions. Graphics were generated with the Aminode tool (*6*).

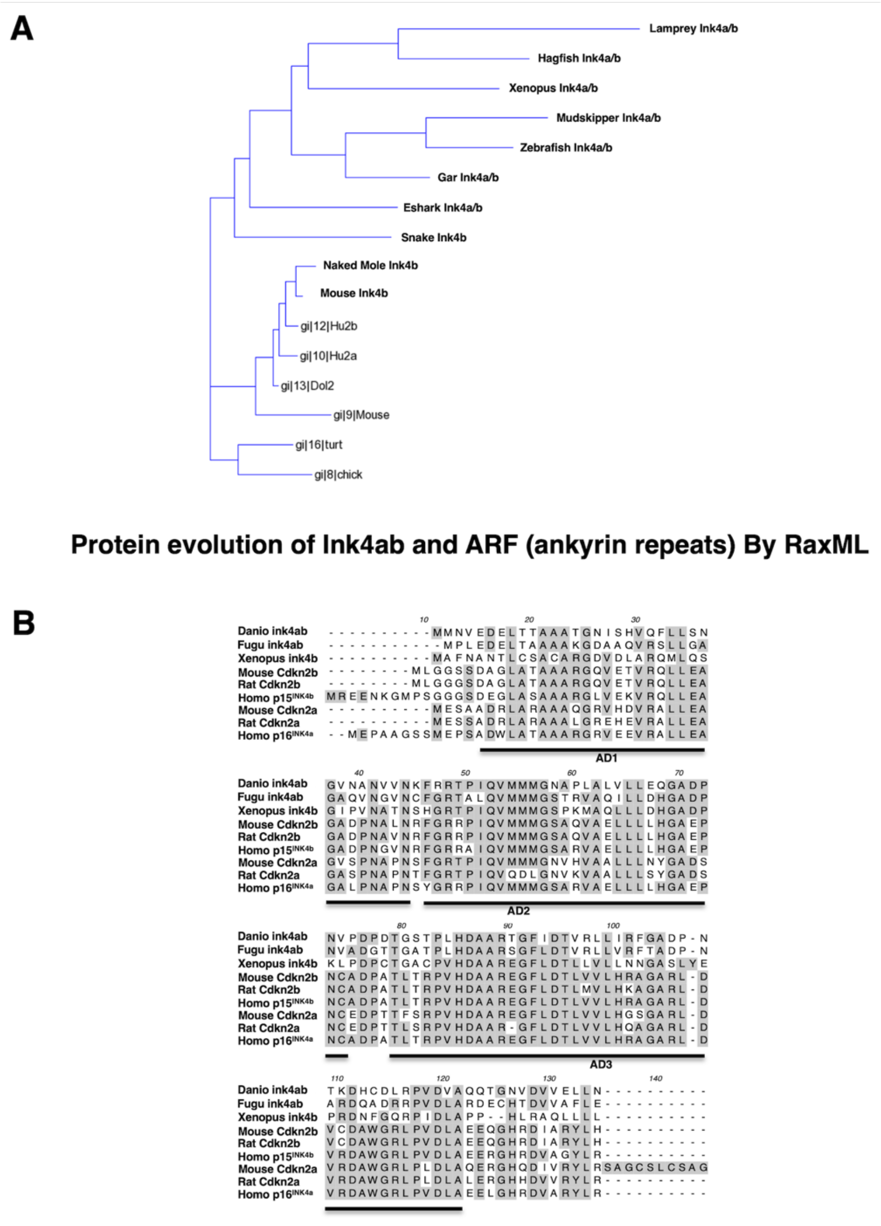


**Fig. S5. Zebrafish ink4ab shares homology with mammalian ink4 proteins at conserved ankyrin repeats.** (**A**) Protein evolutionary tree of Ink4 orthologs. (**B**) ClustalX alignment (*7*) of zebrafish Ink4ab polypeptide sequence with human, mouse, rat, pufferfish (Fugu), and western clawed frog (Xenopus) INK4A and INK4B orthologs. AD, ankyrin domain.

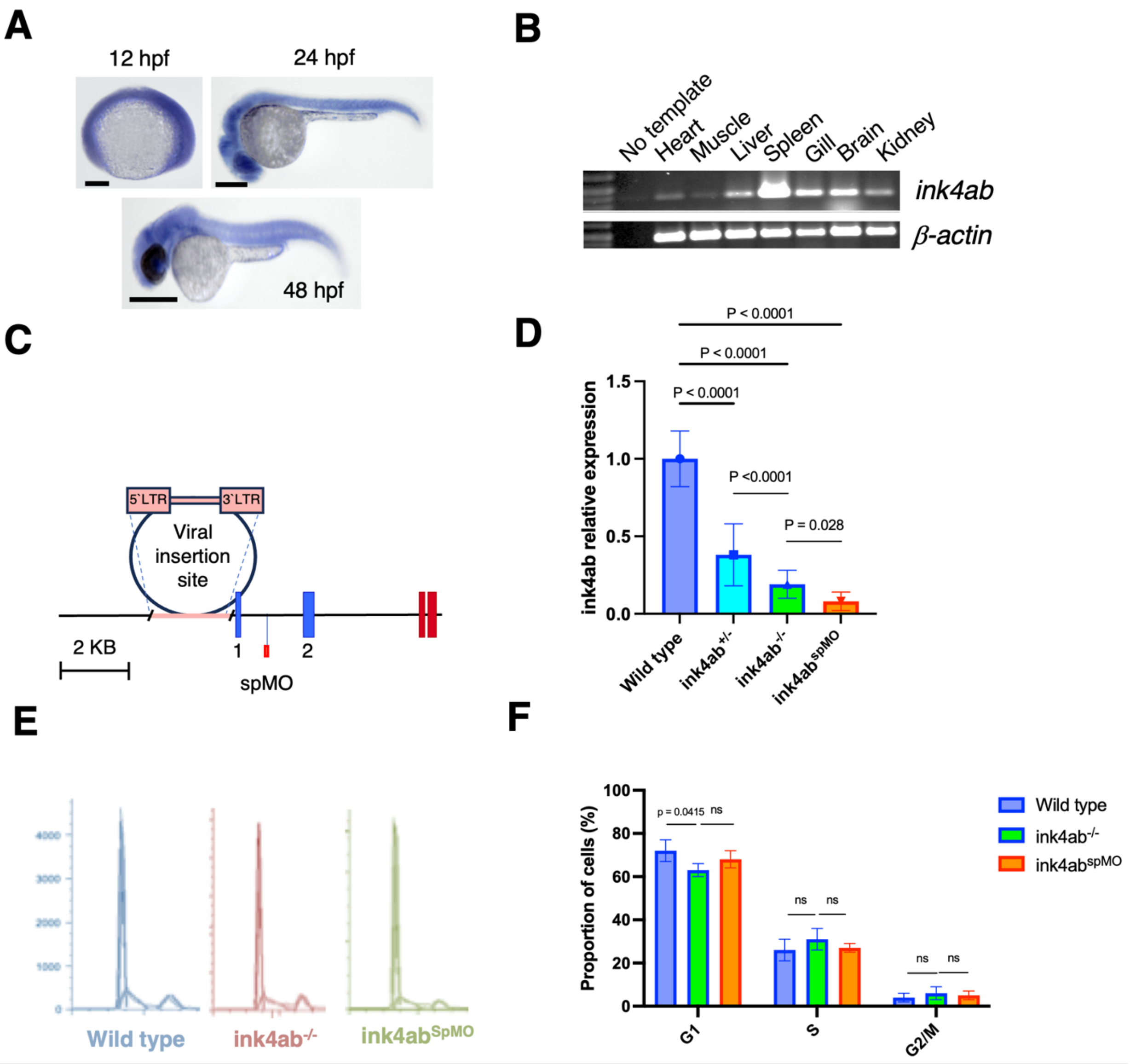


**Fig. S6. Analysis of *ink4ab* expression.** (**A**) ISH to detect *ink4ab* mRNA expression during embryonic development. (**B**) RT-PCR for *ink4ab* expression in adult zebrafish tissues. DNA ladder is on the left, and no template used as a negative control is showing no products. Note the relatively higher expression in the spleen. (**C**) Diagram depicting retroviral integration into the *ink4ab* locus and location of splice morpholinos (spMO).Quantitative analyses of *ink4ab* expression was done using a custom TaqMan Q-RT-PCR assay. (**D**) Confirmed ink4ab deficiency in retroviral mutants and spMO injected embryos using a custom TaqMan Q-RT-PCR assay. (**E**) DNA content profiles of the cell cycle of WT, ink4ab mutants, and spMO injected embryos. (**F**) Proportion of cells in cell cycle phases. Other than a modest but significant reduction in G1 phase of the cell cycle, ink4ab$^{-/-}$ mutant embryos have relatively similar cell cycle profiles compared to WT embryos under normal conditions. hpf, hours post-fertilization. Scale bar is 100 μM.

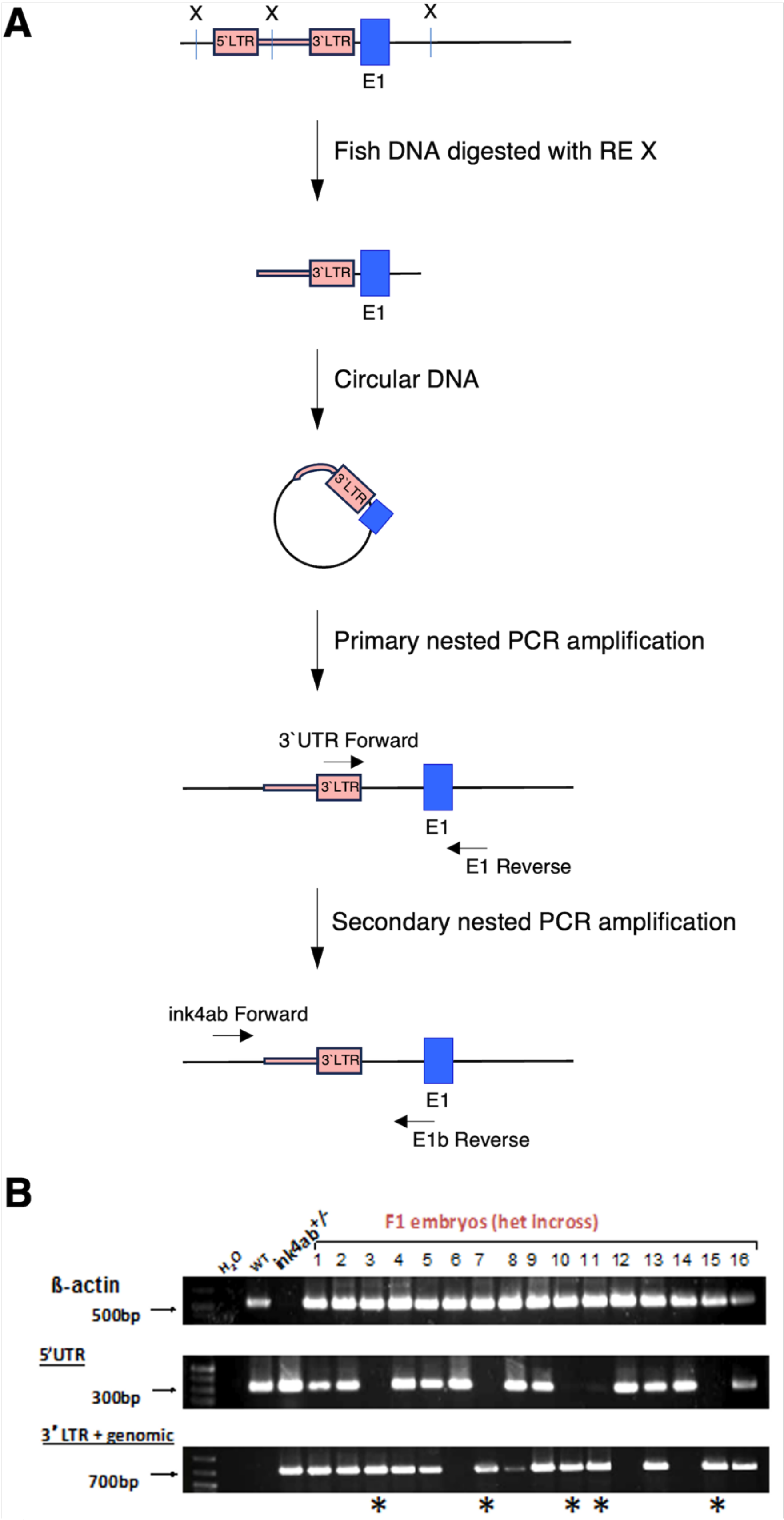
A
X
X
X
5`LTR
3`LTR
E1
Fish DNA digested with RE X
3`LTR
E1
Circular DNA
3`LTR
Primary nested PCR amplification
3`UTR Forward
3`LTR
E1
E1 Reverse
Secondary nested PCR amplification
ink4ab Forward
3`LTR
E1
E1b Reverse
B
F1 embryos (het incross)
H2O
WT
ink4ab+/-
1 2 3 4 5 6 7 8 9 10 11 12 13 14 15 16
ß-actin
500bp
5'UTR
300bp
3' LTR + genomic
700bp
* * * * *

**Fig. S7. Design of retroviral mutagenesis genomic screen to identify *ink4ab* insertional mutant zebrafish.** (**A**) Retroviral insertional mutagenesis was performed to target the zebrafish gene predicted to encode an INK4 gene. Zebrafish were genotyped for verification of the mutation using two nested PCRs. The first reaction included primers specific for viral long terminal repeat sequences (LTRs) to determine that a viral insertion interfering with ink4ab 5`UTR occurred. The second reaction was performed using an aliquot of the first PCR product and primers specific for the zebrafish genomic DNA to amplify the *ink4ab* promoter sequence flanked by the viral LTRs. (**B**) Representative data of genotyping with five female fish and four males were identified as carriers for the viral insertion mutation (asterisks). These fish were used in heterozygous intercrosses to generate the homozygous ink4ab mutants. The gel images are from the nested PCRs.

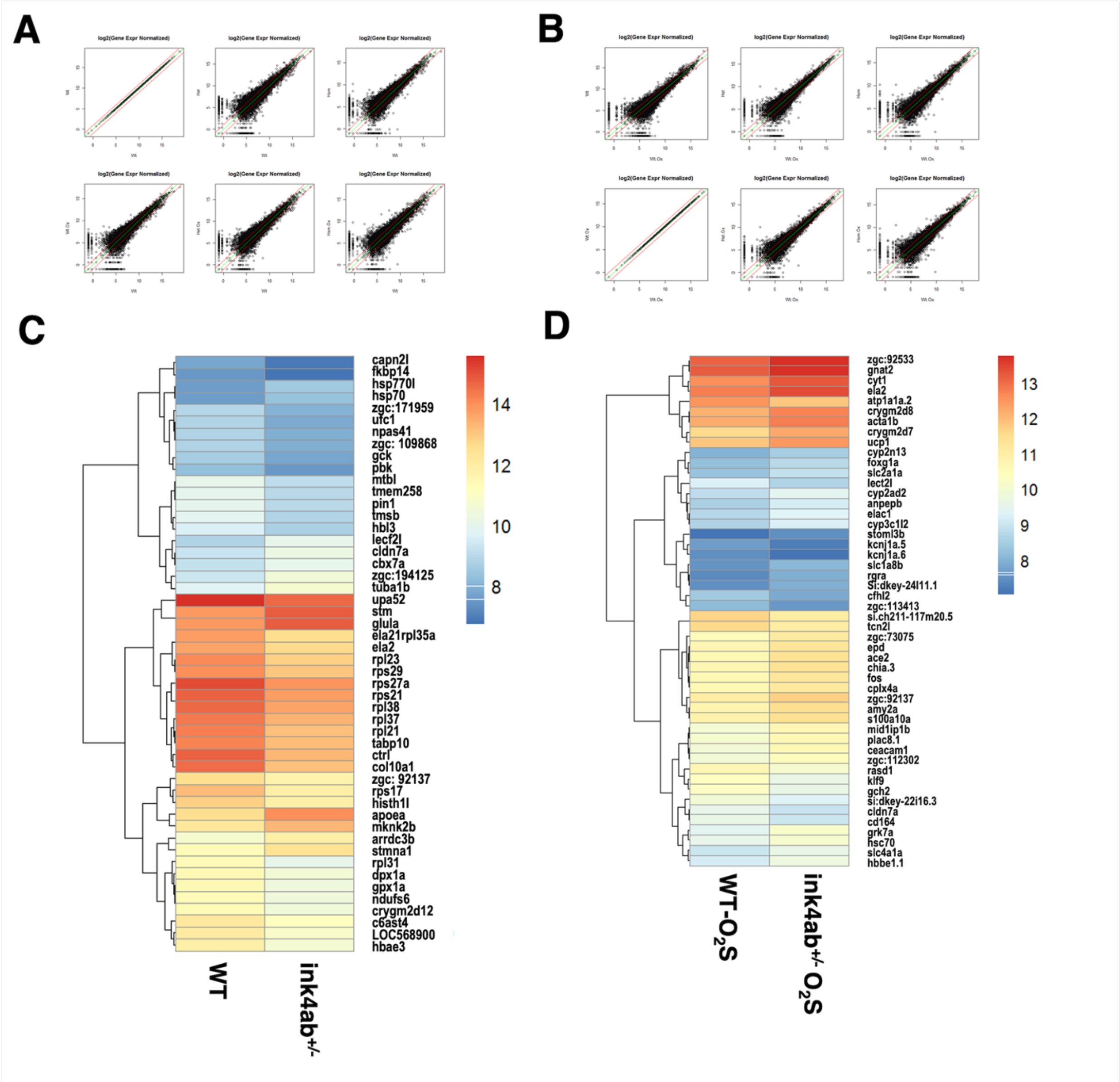


**Fig. S8. RNA transcriptomic analysis of inkab mutant zebrafish under basal and oxidative stress conditions.** (**A-B**) Five-pairwise scatter plots of WT, ink4ab$^{+/-}$ (Het), and ink4ab$^{-/-}$ (Hom) log2 normalized transcriptomic expression showing concordance between the 14,455 gene expression profiles under basal (**A**) or oxidative stress ($O_2S$) (**B**) conditions. (**C-D**) Heatmaps of the top 50 genes which showed transcriptomic changes between WT and Ink4ab$^{+/-}$ embryos to study the effects of single *ink4ab* allele defect under basal (**C**) and $O_2S$ (**D**) conditions from 14,455 transcriptomic targets.

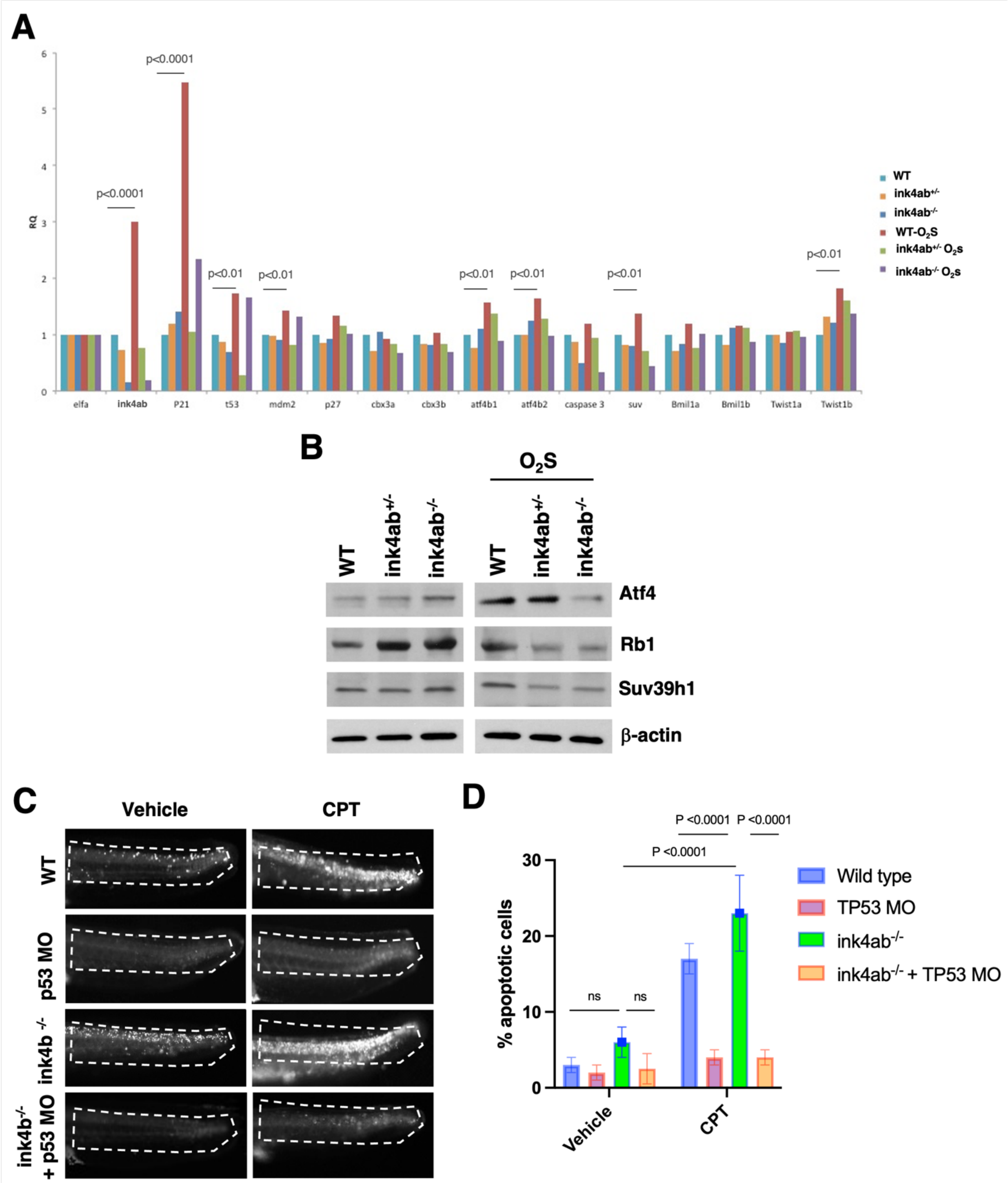


**Fig. S9. P53-dependent apoptosis in inkab mutant zebrafish.** (**A**) Senescence and apoptosis-related transcript changes in WT, Ink4ab$^{+/-}$, and ink4ab$^{-/-}$ under oxidative stress ($O_2S$). Data are shown as normalized transcript levels (RQ) relative to housekeeping (ef1$\alpha$). (**B**) Western Blot of WT, Ink4ab$^{+/-}$, and ink4ab$^{-/-}$ embryo lysates under basal and $O_2S$ conditions. (**C**) Representative acridine orange staining of detect apoptotic cells in vehicle- and CPT-treated embryos (n = 50 embryos/group). (**D**) Quantitation of positive cells in embryo tails.

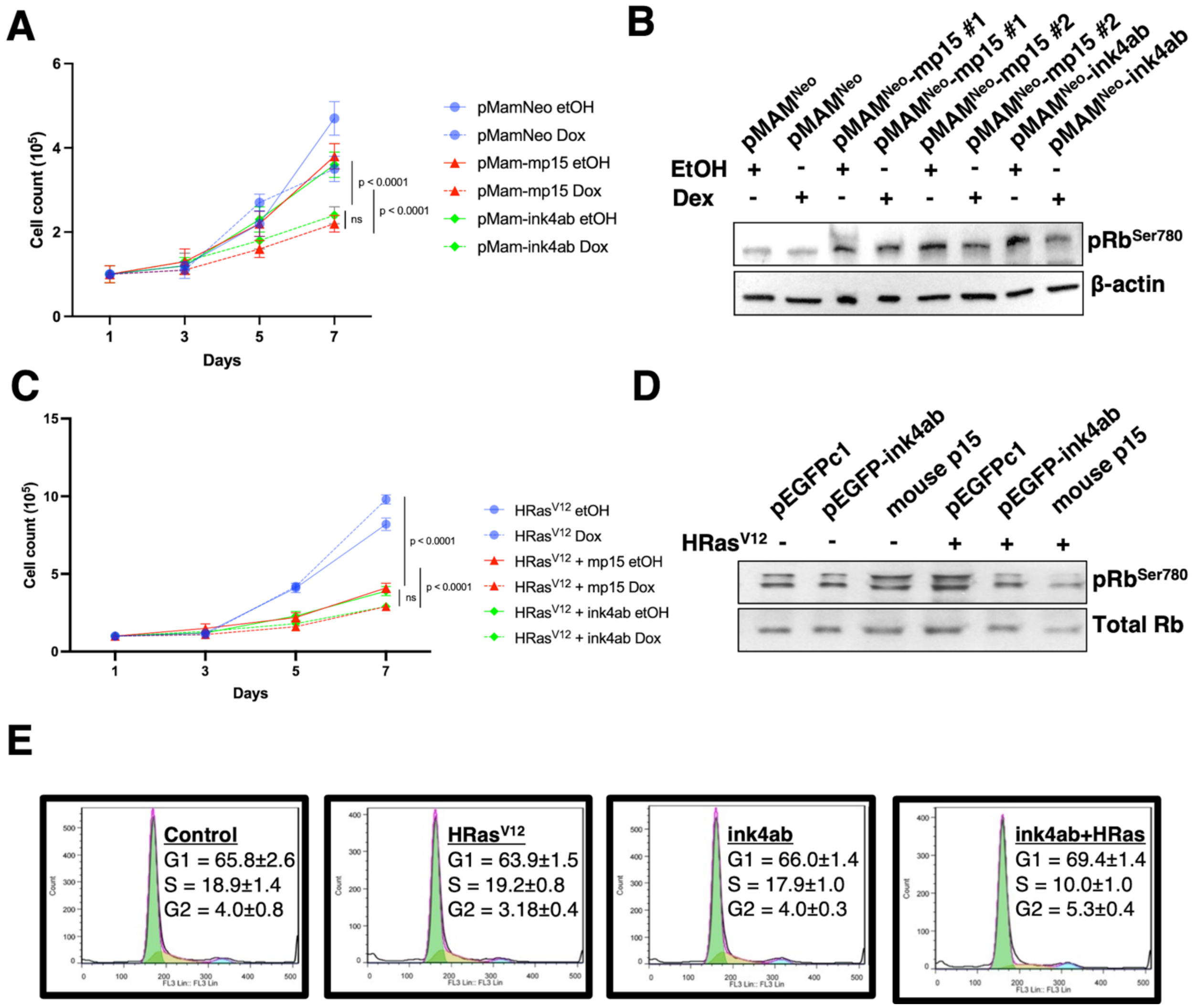


**Fig. S10. Zebrafish ink4ab decreases the growth rate of Ras-transformed mammalian cells.** (**A**) Growth curve of cells transfected with pMam-Neo (vector), pMam-mp15 (mouse p15) or pMam-EGFPink4ab (zebrafish ink4ab) alone and treated with Dexamethasone or ethanol (EtOH). (**B**) Mammalian cells exogenously expressing zebrafish ink4ab show induced expression of phosphor-Rb1 similar to mouse p15. (**C**) Growth of cells co-transfected with pBabeHrasV12 (oncogenic HRas). (**D**) Western blot from lysates of cells transfected with pEGFPc1 (vector), pCR3-mp15 (mouse p15), or pEGFP-*ink4ab* (zebrafish *ink4ab* fused to EGFP). Mammalian cells exogenously expressing zebrafish ink4ab and oncogenic HRas have decreased expression of phosphor-Rb1. (**E**) Cell cycle profiles of 293T cells transfected with either pEGFPc1 or pEGFPink4ab with or without $HRas^{V12}$ and sorted for GFP positive cells. Ink4ab in cells subjected to oncogenic stress with HRas show reduced cells in the G1 phase of the cell cycle. Data are depicted as means ± SD from three independent experiments.

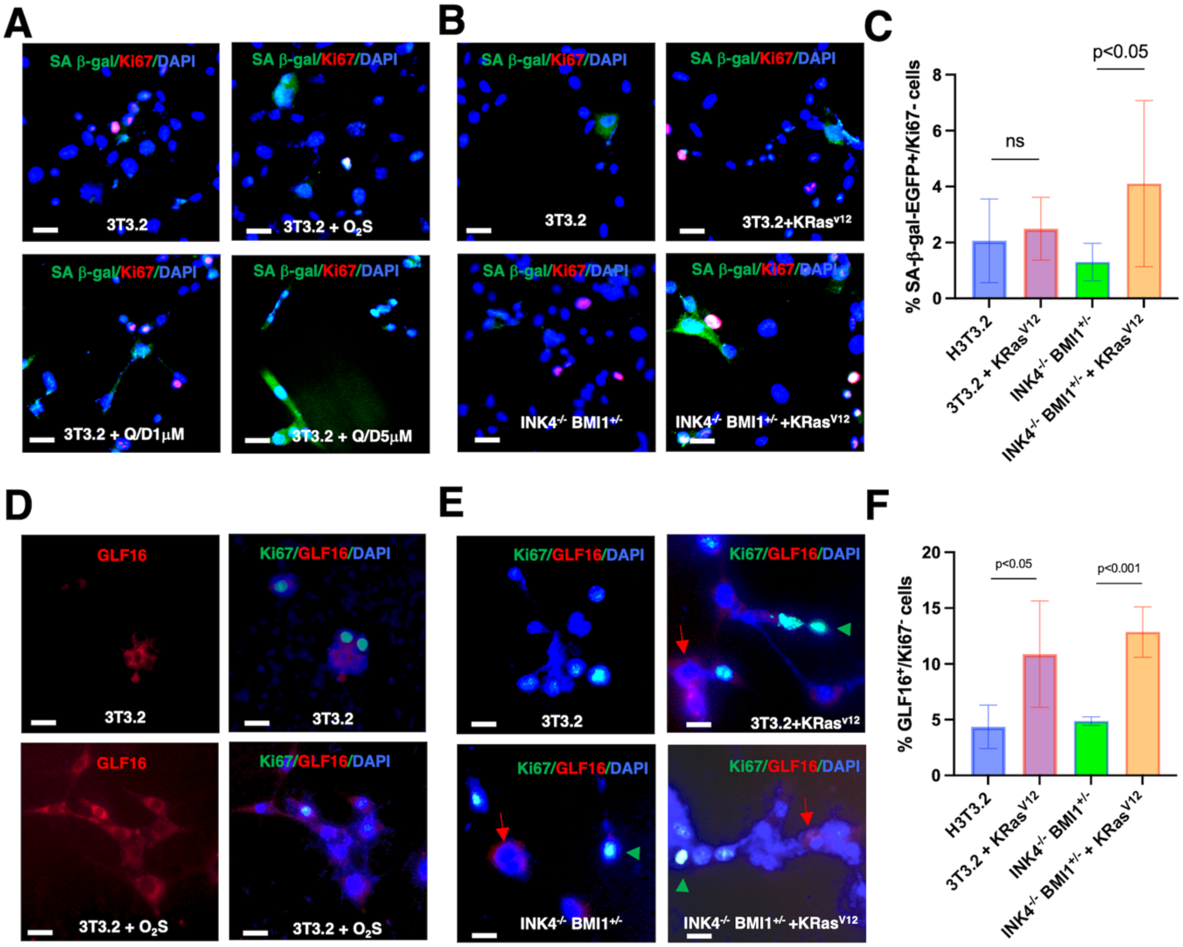


**Fig. S11. Detection of senescent cells by SA-β-gal-EGFP fluorescent probe and validation with GLF16 small molecule. (A-B)** Clone NIH3T3.2 cells and INK4$^{-/-}$/BMI1$^{+/-}$ MEFs transfected with Mock control or KRas$^{v12}$ plasmids were treated or not with vehicle or Quercetin with Dasatinib at 5µM each (Q+D5) for 5 days, then were fixed, permeabilized, and stained with SA-β-gal-EGFP fluorescent probe. Specificity of SA-β-gal-EGFP was evaluated by co-staining with the proliferation marker Ki67. An inverse relationship between SA-β-gal-EGFP and Ki67 positivity was depicted. (**C**) Quantitation of OIS in SA-β-gal-EGFP positive/Ki67 negative senescent NIH3T3.2 and INK4$^{-/-}$/BMI1$^{+/-}$ MEFs with KRas$^{v12}$, depicted as means ± SD from six independent experiments. (**D**) GLF16 (red, alone or with nuclear DAPI) staining in 3T3.2 cells subjected to $O_2S$ resulted in an intense cytoplasmic signal (corresponding to lipofuscin aggregates) abundant in senescent cells relative to untreated cells. (**E**) GLF16 specificity was subsequently evaluated by co-staining with Ki67. An inverse relationship between GLF16 and Ki67 positivity was depicted. (**F**) Quantification of GLF16 positive/Ki67 negative senescent NIH3T3.2 and INK4$^{-/-}$/BMI1$^{+/-}$ MEFs with KRas$^{v12}$ , depicted as means ± SD from six independent experiments. There was an overall concordance between the levels of senescent cells detected with SA-β-gal-EGFP *vs.* GLF16, albeit with a slightly higher background with GLF16 staining. Images were acquired using a confocal microscope equipped with a digital camera and positive cells were evaluated by two independent readers. Scale bars are 20µM.

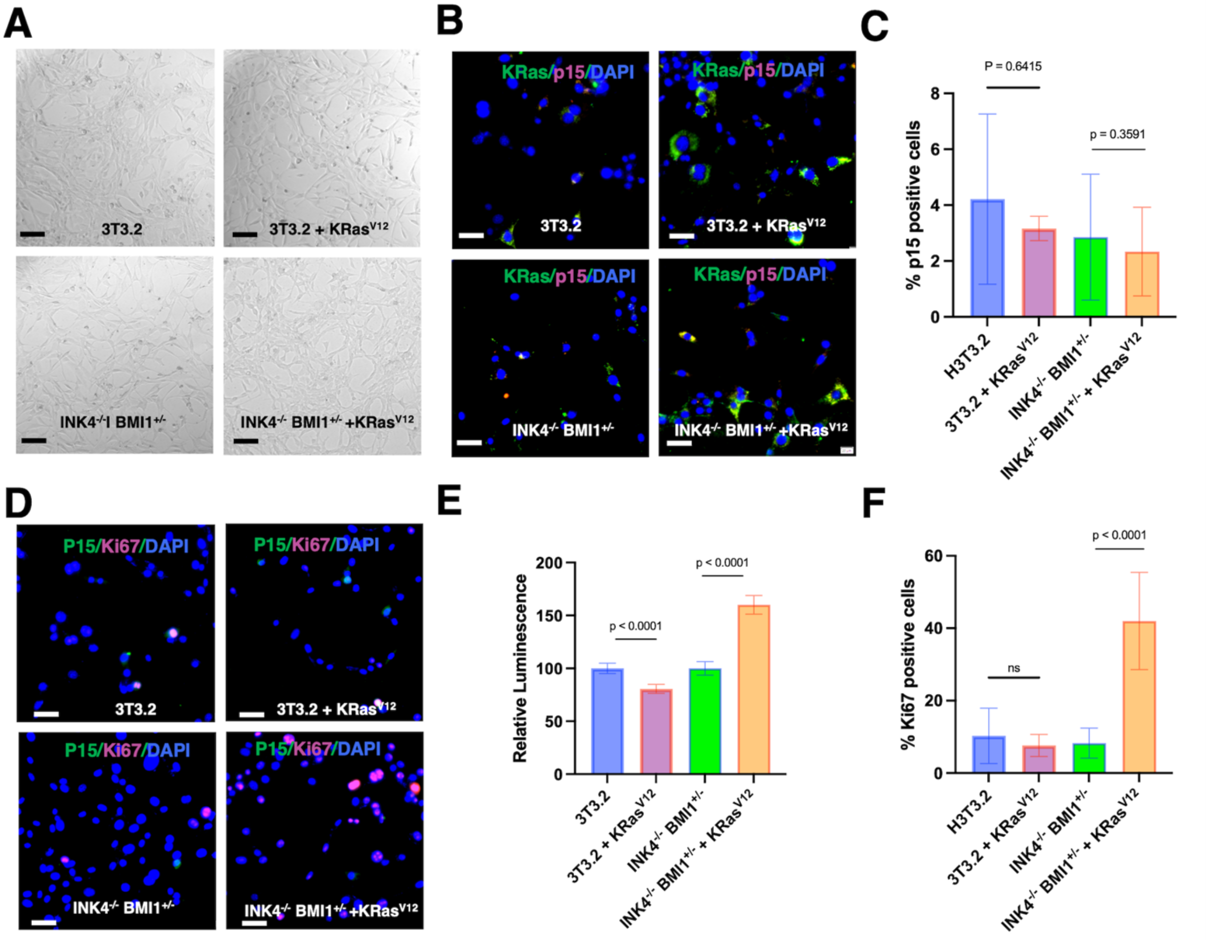


**Fig. S12. Upregulated p15$^{Ink4b}$ in response to oncogenic Kras in MEFs. (A-B)** Clone NIH3T3.2 cells and INK4$^{-/-}$/BMI1$^{+/-}$ MEFs transfected with Mock control or KRas$^{v12}$ plasmids in brightfield images (**A**) and fluorescence images (**B**) showing increased p15$^{Ink4b}$ in response to Kras-mediated OIS. (**C**) Quantitation of increase in p15$^{Ink4b}$ positive cells in response to Kras-mediated OIS, depicted as means ± SD from six independent experiments. (**D**) Fluorescence images showing p15$^{Ink4b}$ in response to Kras-mediated OIS. Note the exclusive staining between p15$^{Ink4b}$ and Ki67 in single cells. (**E**) Reduced proliferation in non-transformed NIH3T3.2 cells, but increased cell proliferation upon loss of INK4 in INK4$^{-/-}$/BMI1$^{+/-}$ MEFs in response to Kras-mediated OIS. (**F**) Quantification of Ki67 positive NIH3T3.2 and INK4$^{-/-}$/BMI1$^{+/-}$ MEFs with KRas$^{v12}$ OIS, depicted as means ± SD from six independent experiments. Images were acquired using a confocal microscope equipped with a digital camera and positive cells were evaluated by two independent readers. Scale bars are 100 μM.

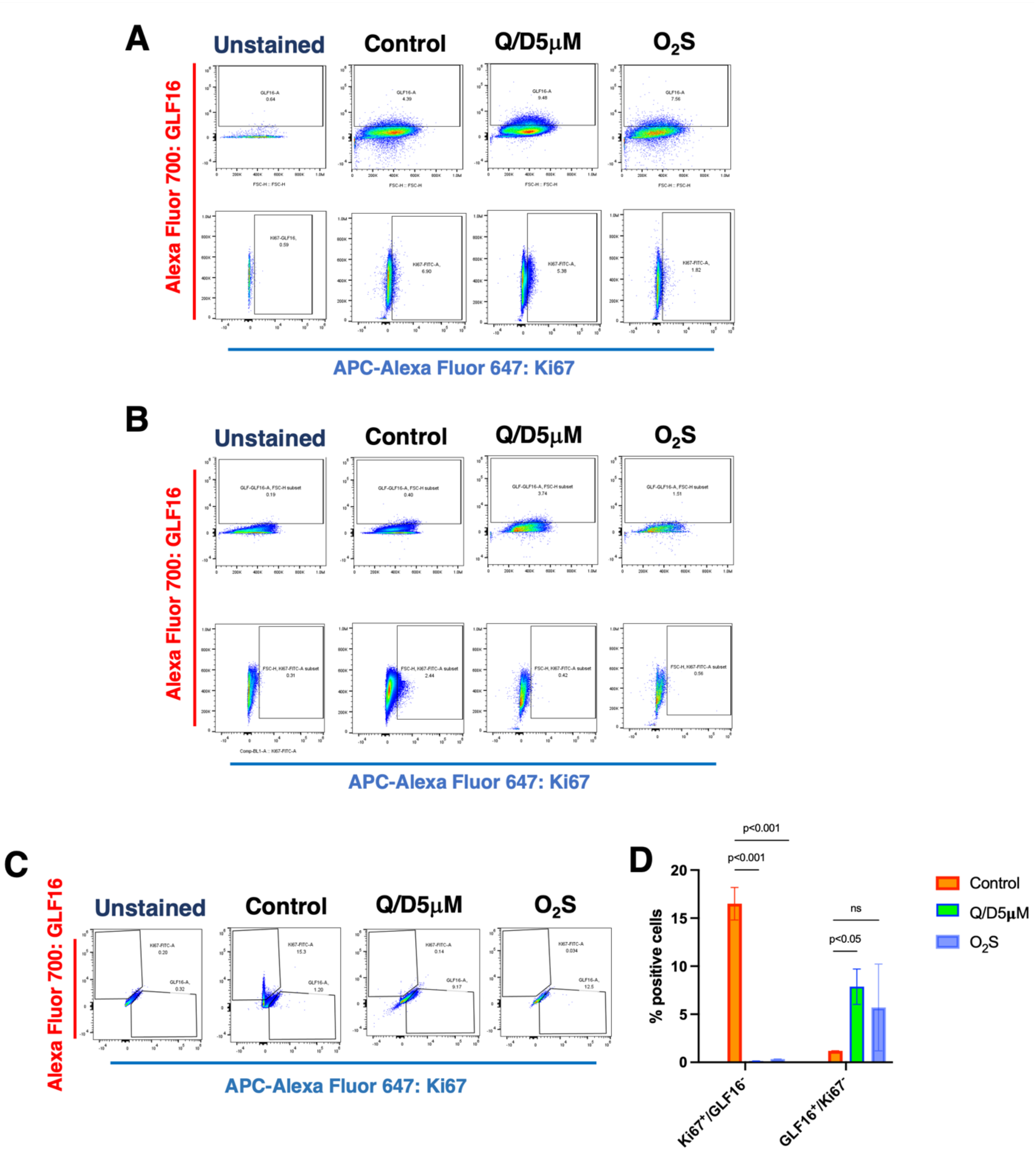


**Fig. S13. Detection of senescent GLF16 positive cells by flow cytometry.** (**A-C**) Proper internalization of GLF16 and specific labeling of senescent cells with $O_2S$ and OIS was validated by flow cytometry in the CRC CT26 cells (**A**), which are deficient in $p15^{Ink4b}/p16^{Ink4a}$ but have $p53^{WT}$ (*28*), and in the CRC MC38 cells (B), which are deficient in $p15^{Ink4b}/p16^{Ink4a}$ and harbor a mutant p53 (*28*). Note that much less senescent cells could be detected and basal GLF16-senescence-associated β-Gal levels were much less in MC38 cells. GLF16 specificity assessed by co-staining of GLF16 with Ki67 in CT26 cells in (**C**). (**D**) Quantitation of positive cells in CT26 and MC38 cells. Data presented as mean ± SD from three independent experiments.

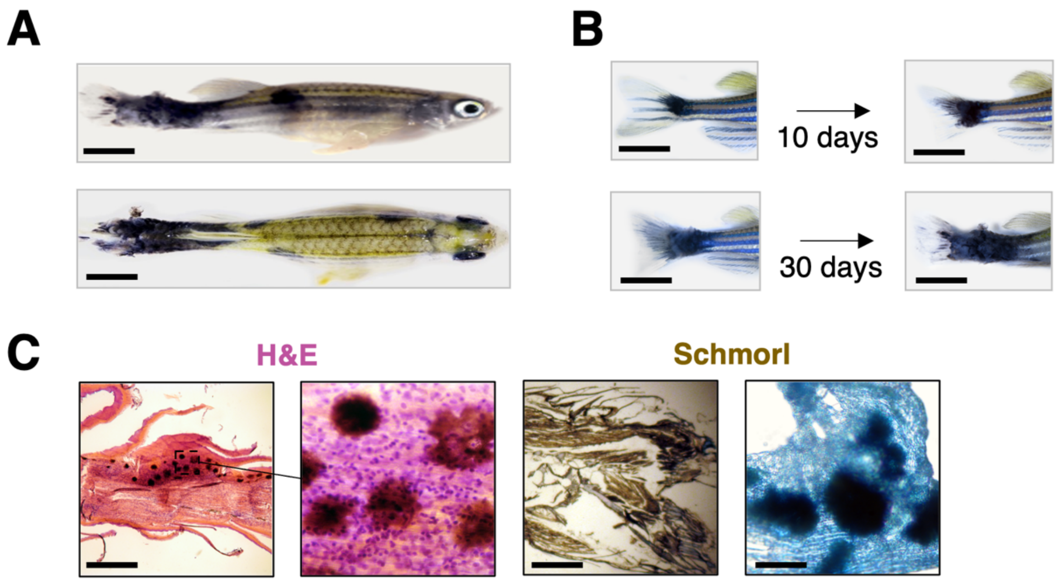


**Fig. S14. Spontaneous melanomas in ink4ab mutant zebrafish.** (**A**) Spontaneous skin melanoma in ink4ab mutant zebrafish were visible more frequently in the fin and tail regions. (**B**) Initial neoplasm observed near tail fin of 5-month-old ink4ab mutant zebrafish, and right image demonstrating the neoplasm spreading and infiltrating tissues at one month later. Scale bars in **A** and **B** are 5mm. **(C)** H&E and Melanocyte-specific Schmorl's stain revealed melanoma metastases throughout an ink4ab mutant fish at one month after initial observation. Scale bar in H&E is 500µM, and the middle image is a higher magnification of the outlined area in the left image. Note the widespread distribution of melanocytic cells in the H&E images. Scale bars in the left and right Schmorl staining images are 200µM and 20µM, respectively. Note that since multiple slides were used in the optimization of staining, the representative Schmorl staining images are from a non-sequential slides from the same melanoma fish presented in **B**, where the melanocytic lesions have spread deeper into the tissue sections.

**A**

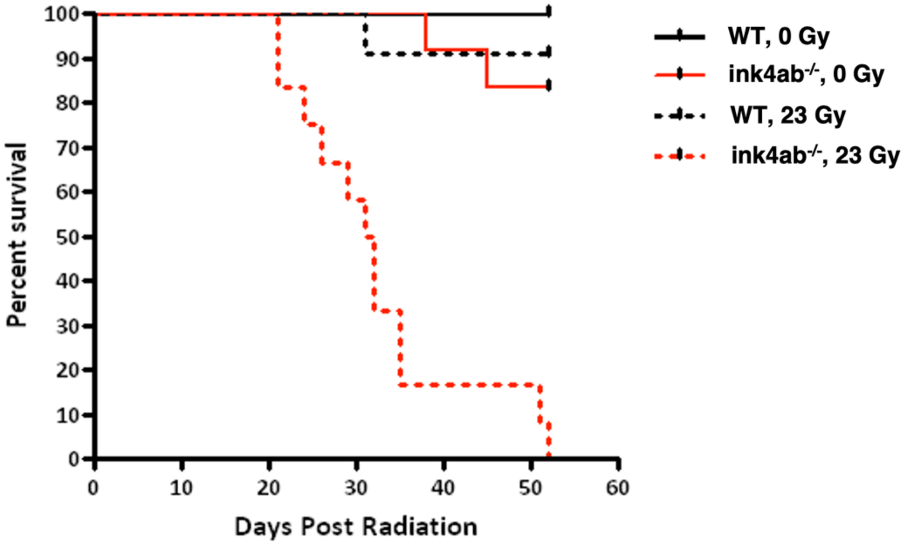


**B**

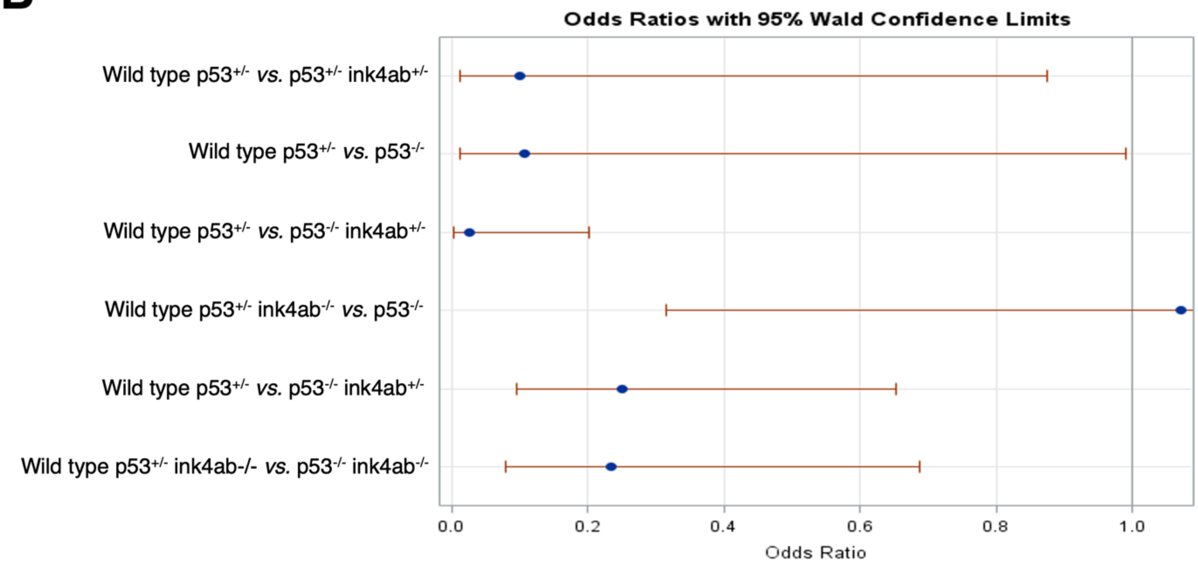


**Fig. S15. Ink4ab deficiency contributes to sensitivity to irradiation and decrease in tumor-free survival of p53 mutant zebrafish. (A)** Kaplan-Meier survival curve of fish subjected to irradiation. The curve shows poor response of ink4ab$^{-/-}$ mutants to sublethal gamma irradiation compared to WT controls. **(B)** Odds Ratios plots on pairwise comparison.

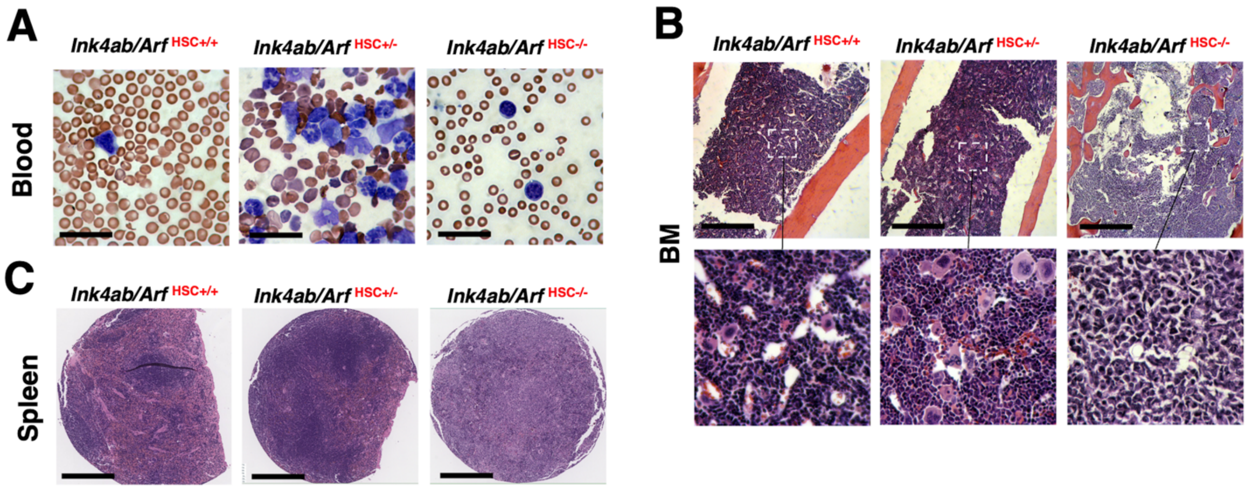


**Fig. S16. Hematopoietic phenotype of conditional ablation of P19$^{Arf}$ in HSCs when combined with p16$^{Ink4A}$ and p15$^{Ink4b}$ deficiency in mice. (A)** Representative images of Giemsa staining of peripheral blood smears from adult 8-month-old mice from either the Ink4ab/Arf$^{HSC+/+}$ cohort, the Ink4ab/Arf$^{HSC+/-}$ cohort, or the Ink4ab/Arf$^{HSC-/-}$ mice. Note that the peripheral blood smears from the Ink4ab/Arf$^{HSC+/-}$ mice frequently displayed an increased number of progenitor-like cells, suggestive of extramedullary hematopoiesis and/or myelodysplastic syndrome-like features. Peripheral blood smears from the Ink4ab/Arf$^{HSC-/-}$ mice presented with anemia and leucopenia with blast-like cells in the peripheral blood. **(B)** Bone marrow (BM) biopsies from the Ink4ab/Arf$^{HSC+/+}$ cohort, the Ink4ab/Arf$^{HSC+/-}$ cohort, or the Ink4ab/Arf$^{HSC-/-}$ mice were stained with H&E, and images were scanned. Note that the BM of the Ink4ab/Arf$^{HSC+/-}$ mice showed increased number of megakaryocytes and progenitor-like cells, suggestive of extramedullary hematopoiesis and/or myelodysplastic syndrome-like features. On the other hand, the BM of the Ink4ab/Arf$^{HSC-/-}$ mice revealed a near complete disappearance of normal or extra-medullary hematopoiesis and BM infiltration with homogenous progenitor-like cells highly suggestive of a leukemia/lymphoma-like phenotype. **(C)** H&E staining of spleens from the three mouse cohorts used for single cell spatial studies. Images were acquired using a brightfield microscope equipped with a digital camera and cell phenotypes were evaluated by two independent readers. Scale bars are 100µM in **A-B** and 200µM in **C**.

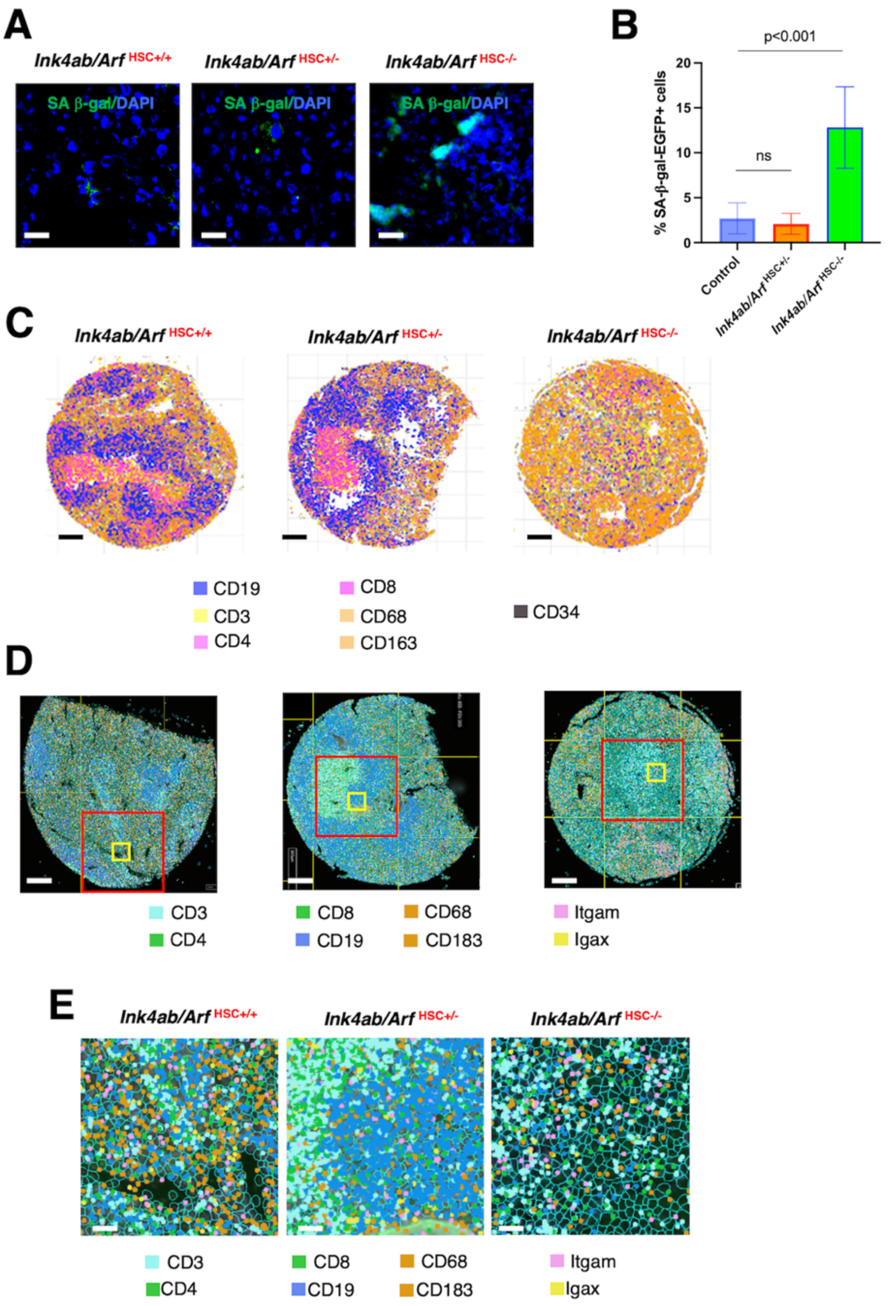


**Fig. S17. Enhanced senescence and impaired HSC differentiation phenotypes upon conditional ablation of P19$^{Arf}$ in HSCs when combined with p16$^{Ink4A}$ and p15$^{Ink4b}$ deficiency in mice. (A)** IF images of mouse liver sections stained with senescence-associated β-Gal-EGFP probe. (**B**) Quantitation of senescence-associated β-Gal-EGFP positive cells. (**C**) Single cell spatial images of mouse splenocytes using CosMx SMI. Cellular neighborhoods are visualized by AtoMx using semi-supervised cell clustering of mouse splenocytes from Ink4ab/Arf$^{HSC+/+}$, Ink4ab/Arf$^{HSC+/-}$, or Ink4ab/Arf$^{HSC-/-}$ mouse spleen sections. Note the single cell spatial loss of CD19+ B-cells and disturbed BCZ and TCZ. (**D**) Overlay of cell types in spatial splenocyte. Red squares outline FOVs used in DEG analyses. (**E**) Cellular neighborhood spatial coordinates of the yellow squared areas in **D**. Scale bars are 100 μM in **A**, **C**, **D**, and 20 μM in **E**.

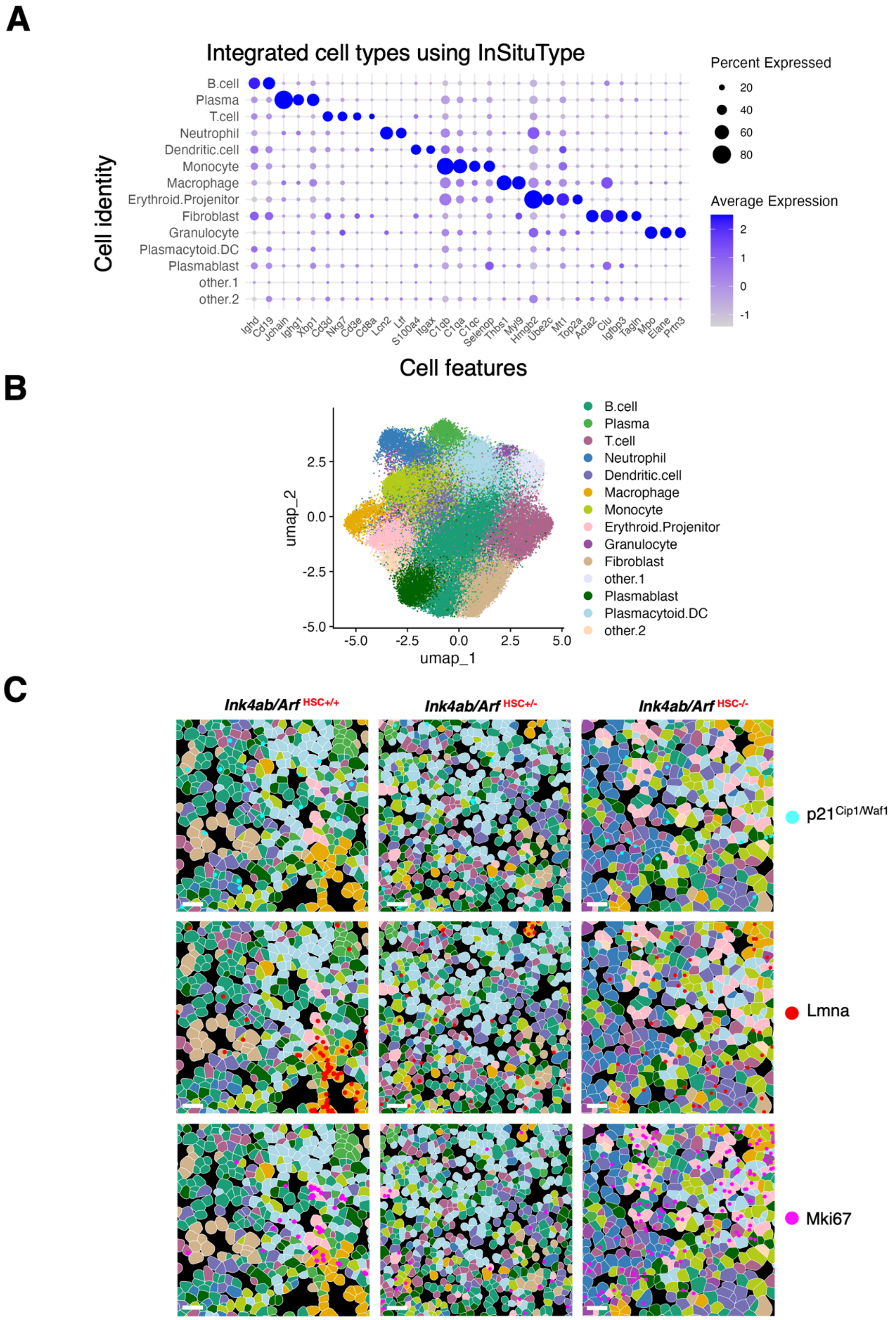
A
Integrated cell types using InSituType
Cell identity
B.cell
Plasma
T.cell
Neutrophil
Dendritic.cell
Monocyte
Macrophage
Erythroid.Projenitor
Fibroblast
Granulocyte
Plasmacytoid.DC
Plasmablast
other.1
other.2
Ighd Cd19 Jchain Ighg1 Xbp1 Cd3d Nkg7 Cd3e Cd8a Lcn2 Ltf S100a4 Itgax C1qb C1qa C1qc Selenop Thbs1 Myl9 Hmgb2 Ube2c Mt1 Top2a Acta2 Clu Igfbp3 Tagln Mpo Elane Prtn3
Cell features
Percent Expressed
20
40
60
80
Average Expression
2
1
0
-1
B
umap_2
umap_1
2.5
0.0
-2.5
-5.0
-5.0
-2.5
0.0
2.5
5.0
B.cell
Plasma
T.cell
Neutrophil
Dendritic.cell
Macrophage
Monocyte
Erythroid.Projenitor
Granulocyte
Fibroblast
other.1
Plasmablast
Plasmacytoid.DC
other.2
C
Ink4ab/Arf HSC+/+
Ink4ab/Arf HSC+/-
Ink4ab/Arf HSC-/-
p21Cip1/Waf1
Lmna
Mki67

**Fig. S18. Single cell spatial deconvolution of splenocytes. (A)** A heatmap plot displaying the estimated mean cell type proportion for each cell type in each spatial splenocyte domain. Processed transcriptomic and proteomic datasets were analyzed with InSituType custom scripts for visualization, cell-state annotation confirmed with FindMarkers, metacell inference, and multimodal integration, and Seurat v5.0.1 package was used for cell typing The color scale is normalized to the -1 to 2 range. Point size indicates the estimated percentage of cell types expressed. (**B**) Single cell spatial analyses of splenocyte phenotypes. Cellular neighborhoods in CosMx data visualized by Uniform Manifold Approximation and Projection (UMAP) using unsupervised cell clustering of mouse splenocytes from Ink4ab/Arf$^{HSC+/+}$, the Ink4ab/Arf$^{HSC+/-}$, and Ink4ab/Arf$^{HSC-/-}$ mouse spleen sections. **(C)** Spatial domain distribution of splenic microenvironment clusters. High-resolution spatial capture (0.2 μm/pixel) across FOVs were processed for each tissues, filtered with a custom pipeline, and log library size normalized. Images represent single molecule display of the same FOV multichannel overlay of senescence-related targets p21$^{Cip1/Waf1}$ (neon blue), nuclear lamin (red), and proliferation signal Mki67 (pink) from Main manuscript **Fig 5H**. Cell type color indicates clusters from **(B)**. Round point color represent senescence associated p21$^{Cip1/Waf1}$ (neon blue), nuclear lamin-a (red), and proliferation signal Mki67 (pink). Note that the individual senescence-associated markers in these same FOVs are the same as the FOVs in **Fig. 5H** and were used to confirm the specificity of p21$^{Cip1/Waf1}$, Lamin, and Mki67 staining in single splenocytes. A mutually exclusive relationship between p21$^{Cip1/Waf1}$ and Mki67 positivity was depicted. Scale bars are 20 μM.

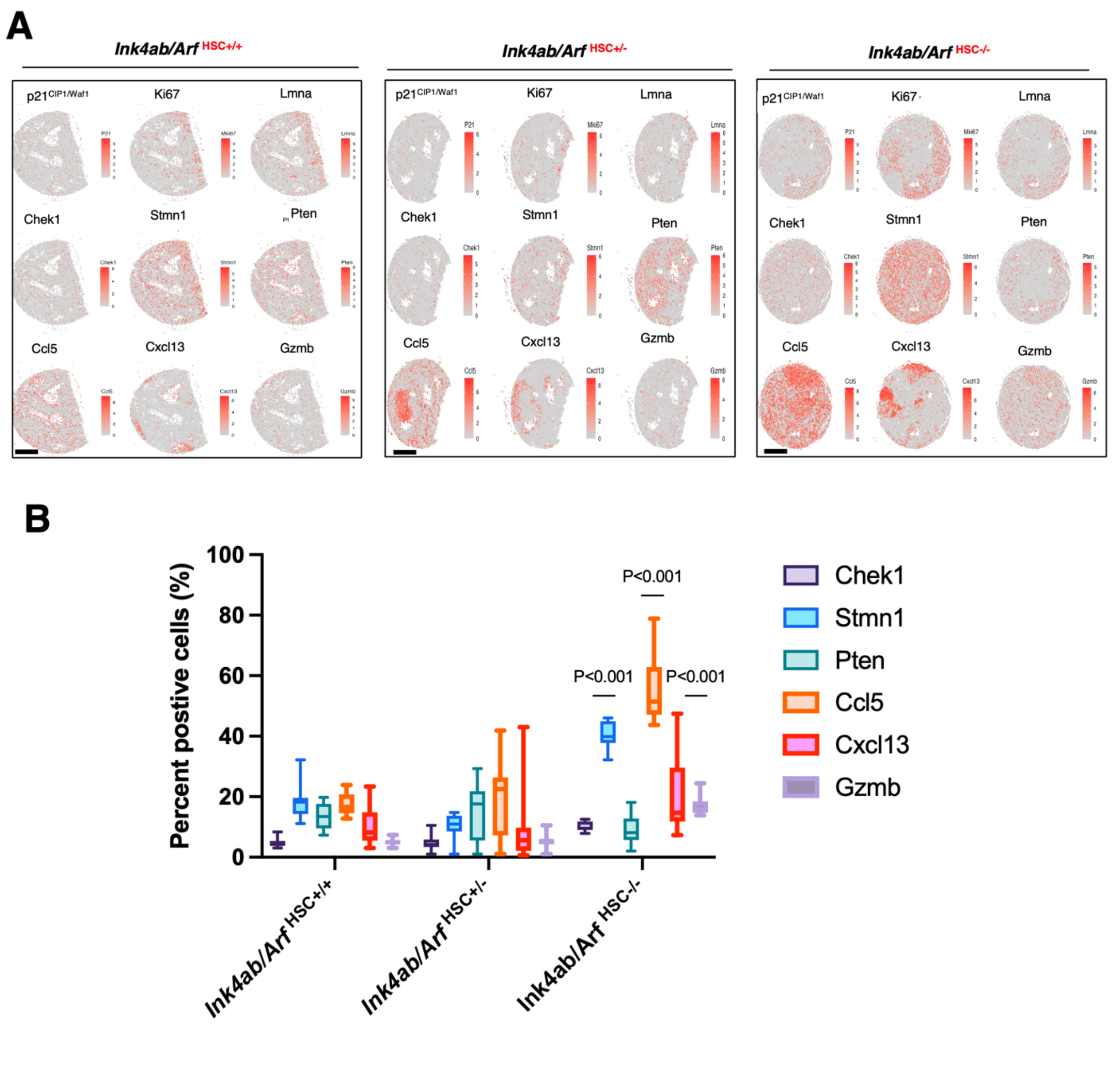


**Fig. S19. Senescence signature of single splenocytes with spatial distribution. (A)** Scatter plot displaying the spatial distribution of single-cell expression profiles across 9 key senescence-associated markers encoded in the splenocyte sections. Senescence markers were selected to represent various types of senescence, including (1) Cell cycle inhibitors: activated p21$^{Cip1/Waf1}$, lack of Mki67 proliferation marker in the same single cell, and reduced nuclear lamins, correlating with senescence-derived epigenetic spatial rearrangement of H3K9me3 to form senescence-associated hetero-chromatin foci (SAHF) (see **fig. S21**), (2) DNA damage repair (DDR) and deregulated metabolic senescence: activated Chek1 for senescent persistent nuclear DNA damage foci called DNA segments with chromatin alterations reinforcing senescence (DNA SCARSs), and activated Stmn1 and reduced PTEN for deregulated metabolic profiles, and (3) SASP: activated Ccl5, Cxcl13, and Gzmb for splenocyte-associated SASP. Single cell spatial profiles of each senescence-associated target is colored according to log2 transformation of the sum of pixel intensities in each cell, with the threshold for visualization being the geometric mean of negative isotype controls (mouse IgG1, mouse IgG2a, rabbit IgG) plus 1.5 standard deviations. (**B**) Percentage of positive cells expressing deregulated metabolic markers and SASP cytokines Ccl5 and Cxcl13 were significantly upregulated in splenocytes from Ink4ab/Arf$^{HSC-/-}$ mice relative to splenocytes from the Ink4ab/Arf$^{HSC+/+}$ mice. Data are depicted as means ± SD from 14 FOVs. Scale bars are 500 μM.

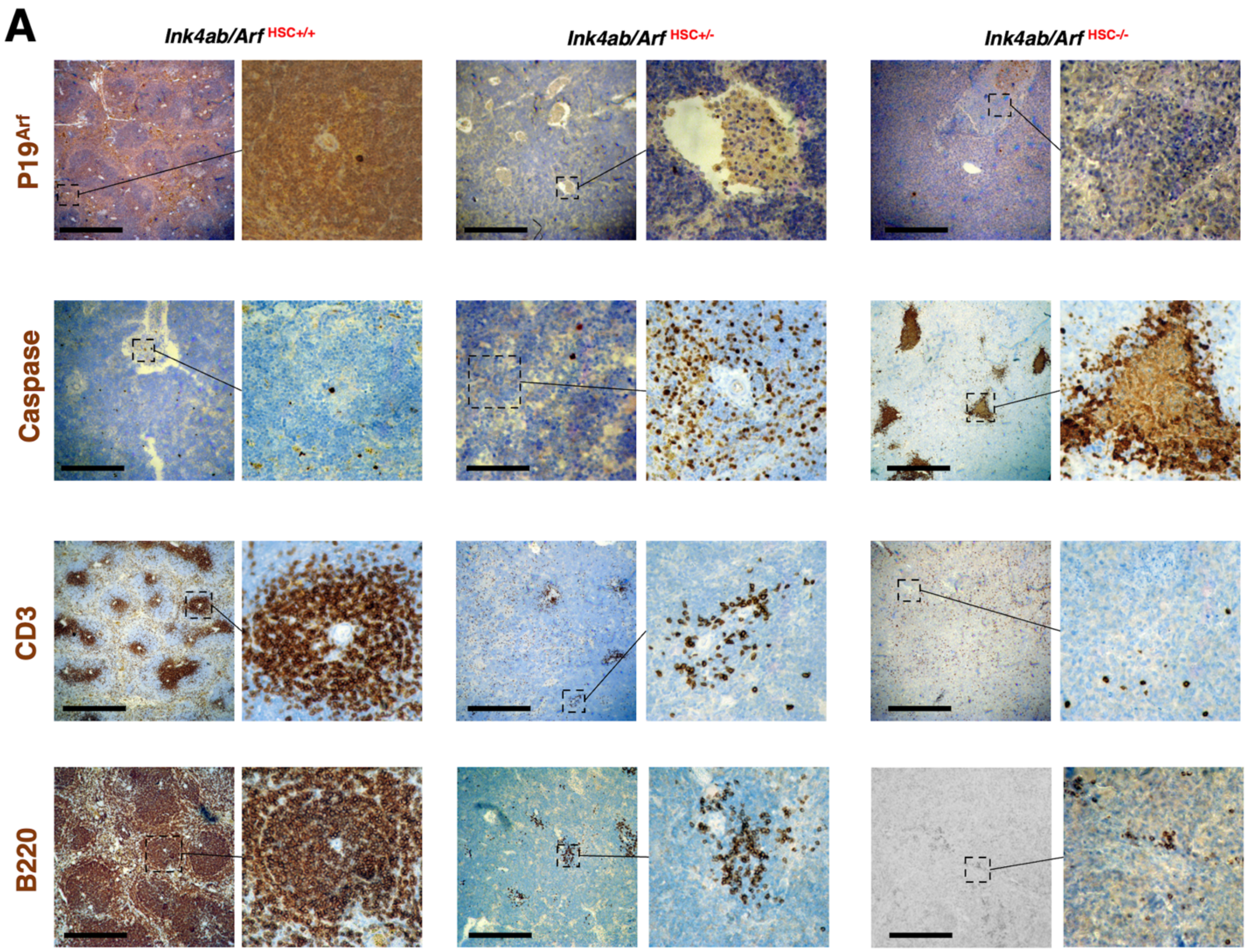


**Fig. S20. Immunohistochemical (IHC) validation of splenic cell phenotypes. (A)** Representative images from IHC staining of Ink4ab/Arf$^{HSC+/+}$, the Ink4ab/Arf$^{HSC+/-}$, and Ink4ab/Arf$^{HSC-/-}$ mouse spleen sections. IHC with the anti-p19$^{Arf}$ antibody was used to confirm the conditional loss of p19$^{Arf}$ in Ink4ab/Arf$^{HSC-/-}$ mouse spleens relative to Ink4ab/Arf$^{HSC+/+}$. Activated caspase-3 antibody was utilized to demonstrate active apoptosis in the T-cell zone (TCZ), which is also called the periarteriolar lymphoid sheath (PALS) because it forms around the central arteriole (outlined area) that runs through the white pulp (WP) on its way to the red pulp (RP)-WP border. T-cells and B-cells was validated with anti-CD3 and anti-B220 antibodies, respectively. Images on the right are higher magnifications of the outlined areas in the images on the left. Scale bars are 200 μM.

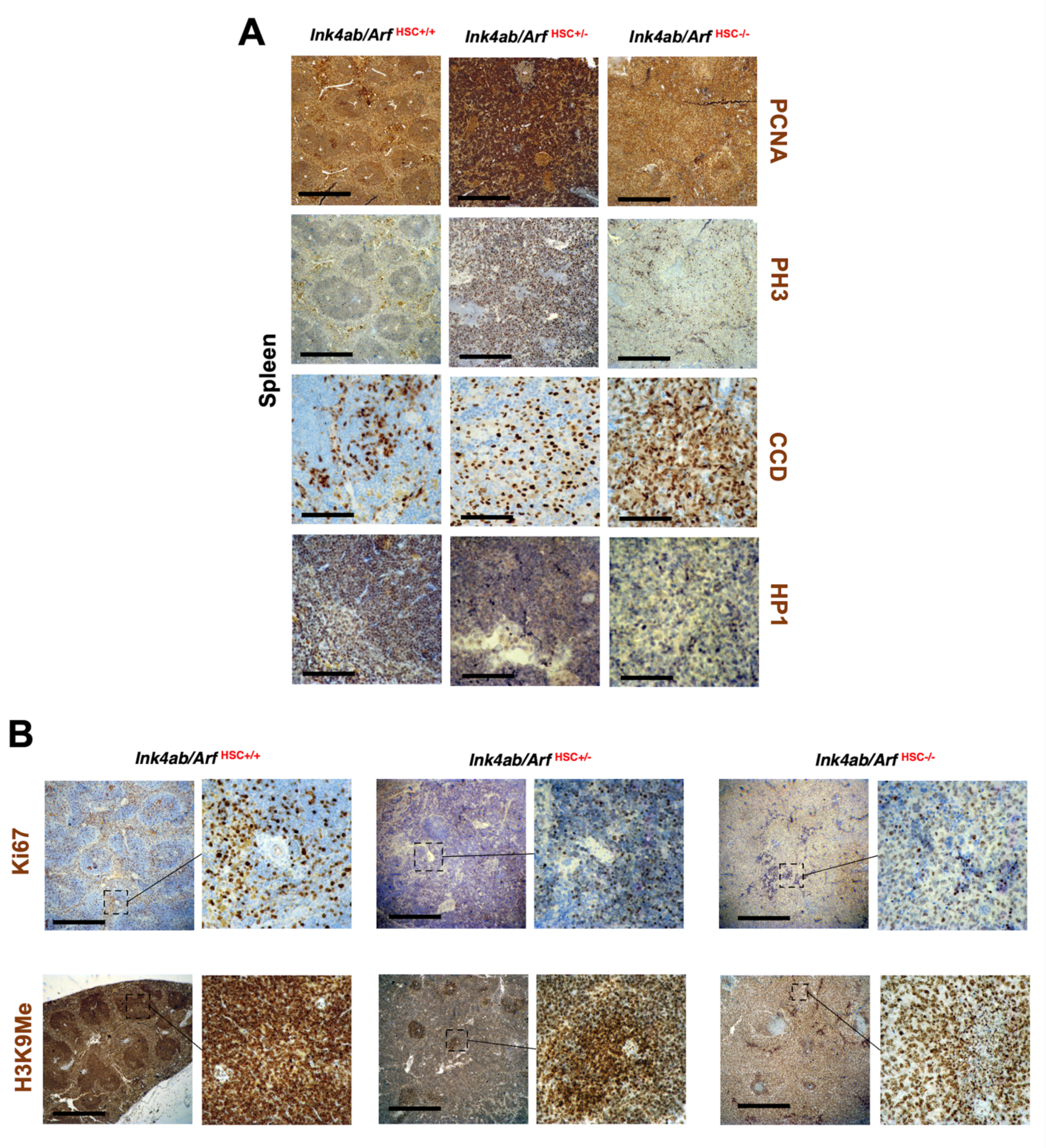


**Fig. S21. IHC analyses of senescence and proliferation markers in splenocytes. (A)** Representative images from IHC staining of Ink4ab/Arf$^{HSC+/+}$, the Ink4ab/Arf$^{HSC+/-}$, and Ink4ab/Arf$^{HSC-/-}$ mouse spleen sections for proliferation marker (PCNA), cell cycle mitosis (PH3), cyclin D1 (CDD), and Heterochromatin-1 (HP1) correlating with SAHF. (**B**) Reduced ki67 expressing cells in the TCZ or PALS around the central arteriole, which is the site for activated T-cells at the RP-WP border, and senescence-derived epigenetic spatial rearrangement of H3K9me3 to form SAHF. Images on the right are higher magnifications of the outlined areas in the images on the left. Scale bars are 200 μM.

**Table S1. The overall 2-year survival of *ink4ab* mutant zebrafish.** Top, statistics of total fish censored and death events. Middle, overall log rank test comparing survival between groups. Matched controls from the same colonies were also manipulated in the same way, including anesthetization, but were not subjected to radiation. All groups were then recovered from anesthesia and monitored for survival. These fish were censored until the last ink4ab mutant fish had died at 52 days post radiation. We observed 12 fish per group daily for survival. We also employed a Cox proportional hazard model on the data using non-irradiated ink4ab$^{-/-}$ as a reference group. This test shows that overall risks of dying for fish both WT and ink4ab$^{-/-}$ exposed to radiation is different from one for fish with ink4ab$^{-/-}$ without radiation. Bottom, Cox proportional hazard ratios for *ink4ab* mutants compared to WT.

| Group | Total (n) | Death (n) | Censored (at 24 M) | Censored (%) |
|---|---|---|---|---|
| Group 1 (ink4ab$^{+/-}$) | 27 | 11 | 16 | 59.26 |
| Group 2 (ink4ab$^{-/-}$) | 161 | 110 | 51 | 31.68 |
| Group 1 (WT) | 124 | 18 | 106 | 85.48 |
| Total | 312 | 139 | 173 | 55.45 |

| Comparison | $X^2$ Test | P-Value |
|---|---|---|
| ink4ab$^{+/-}$ *vs.* ink4ab$^{-/-}$ | 59.67 | P < 0.001 |
| ink4ab$^{+/-}$ *vs.* WT | 46.49 | P < 0.001 |
| ink4ab$^{-/-}$ *vs.* WT | 98.23 | P < 0.001 |

| | Estimate (S.E) | Hazard Ratio (HR) | 95% CI for HR | P-value |
|---|---|---|---|---|
| Group (ink4ab$^{+/-}$) | 1.27 (0.38) | 3.58 | [1.69, 7.57] | P < 0.001 |
| Group (ink4ab$^{-/-}$) | 2.16 (0.26) | 8.64 | [5.21, 14.33] | P < 0.001 |

**Table S2. Survival analysis of Gamma-irradiated ink4ab fish.** Median survival of wild type and *ink4ab* mutants subjected to radiation. Overall log rank test comparing survival between groups. Cox proportional hazard ratios for irradiated wild type and ink4ab$^{-/-}$ fish compared to non-irradiated *ink4ab*$^{-/-}$. overall survival time between groups, the Sidak multiple-comparison adjustment was used. There is significant difference in survival times in *ink4ab* without radiation vs. *ink4ab* with radiation and WT with radiation vs. ink4ab with radiation, but not in *ink4ab* without radiation *vs.* WT with radiation. We also employed a Cox proportional hazard model on the data using non-irradiated ink4ab$^{-/-}$ as a reference group. This test shows that overall risks of dying for fish both WT and ink4ab$^{-/-}$ exposed to radiation is different from one for fish with *ink4ab*$^{-/-}$ without radiation.

| Comparison | $X^2$ Test | P-Value |
|---|---|---|
| ink4ab No Rad *vs.* WT + Rad | 0.011 | P = 0.9964 |
| ink4ab No Rad *vs.* ink4ab + Rad | 21.54 | P < 0.001 |
| WT + Rad *vs.* ink4ab + Rad | 23.8 | P < 0.001 |

| | Estimate (S.E) | Hazard Ratio (HR) | 95% CI for HR | P-value |
|---|---|---|---|---|
| Group (WT + Rad) | -0.6 (1.22) | 0.551 | [0.05, 5.997] | P = 0.627 |
| Group (ink4ab$^{-/-}$ + Rad) | 2.79 (0.79) | 16.275 | [3.461, 76.585] | P = 0.0004 |

**Table S3. Ink4ab deficiency contributes to a decrease in tumor-free survival of p53 mutant zebrafish.** Inference results for odds ratios (OR) using $p53^{+/-}$ as a reference, and OR estimates on pairwise comparison.

| | Estimate (S.E.) | OR (95% C.I.) | P-value |
|---|---|---|---|
| **Intercept** | -2.77 (1.03) | | 0.0071 |
| **$p53^{-/-}$** | 2.3 (1.11) | 9.99 [1.14, 87.42] | 0.0375 |
| **$p53^{+/-}$ $ink4ab^{+/-}$** | 2.23 (1.14) | 9.33 [1.01, 86.29] | 0.0491 |
| **$p53^{-/-}$ $ink4ab^{-/-}$** | 3.69 (1.07) | 39.89 [4.93, 323.98] | 0.0006 |

| Comparison | OR | 95% C.I. |
|---|---|---|
| **$p53^{+/-}$ *vs.* $p53^{+/-}$ $ink4ab^{-/-}$** | 0.1 | [0.011, 0.875] |
| **$p53^{+/-}$ *vs.* $p53^{-/-}$** | 0.107 | [0.012, 0.992] |
| **$p53^{+/-}$ *vs.* $p53^{-/-}$ $Ink4ab^{-/-}$** | 0.025 | [0.003, 0.203] |
| **$p53^{+/-}$ $Ink4ab^{+/-}$ *vs.* $p53^{-/-}$** | 1.071 | [0.316, 3.636] |
| **$p53^{+/-}$ $Ink4ab^{+/-}$ *vs.* $p53^{-/-}$ $Ink4ab^{-/-}$** | 0.25 | [0.096, 0.653] |
| **$p53^{-/-}$ *vs.* $p53^{-/-}$ $Ink4ab^{-/-}$** | 0.233 | [0.079, 0.687] |

**Table S4. CosMx single cell spatial FOV summary.**

| | *Ink4ab/Arf* HSC+/+ | *Ink4ab/Arf* HSC+/- | *Ink4ab/Arf* HSC-/- |
|---|---|---|---|
| **Number of FOVs** | 14 | 13 | 18 |
| **Total tissue area ($mm^2$)** | 8.06 | 8.02 | 8.86 |
| **Mean single cell size ($\mu m^2$)** | 61.9 | 62.3 | 64.4 |
| **Number of cells in final analysis** | 26,440 | 12,582 | 35,768 |
| **Total transcripts assigned to cells** | 6,838,950 | 3,314,468 | 5,540,229 |
| **Mean transcripts per cell** | 259 | 263 | 155 |
| **Mean transcript per $\mu m^2$** | 4.2 | 4.23 | 2.4 |

**Table S5. Spatial DEGs.**

**See the attached data file.**

**Table S6. Primer sequences for genotyping of ink4ab zebrafish line.**

| Primer | **Sequence (5'-3')** |
|---|---|
| **3'LTR (viral)** | CACCAGCTGAAGCCTATAGAGTACGAGC |
| 150467-3-92 (GENOMIC) | TCGCTTCACAGGTAAAACCAA |
| **3'LTR NEST 1 (VIRAL)** | AGTCTCCAGAAAAAGGGGGGAATG |
| 150467-3-86 (GENOMIC) | CACAGGTAAAACCAAATGATG |
| p**16VIfor** | GCAGTATGACAGTACTGCACC |
| p16VIrev | GGGAAATGTTTCCGGTGGC |

**Table S7. Antibodies and conditions used for western blotting**.

| Antibody | **Company** | **Dilution** |
|---|---|---|
| p15 (K-18) | Santa Cruz | 1:2,000 |
| β-actin (AC-15) | Santa Cruz | 1:500 |
| Phospho-RB1(Ser 780) | Cell signaling | 1:1,000 |
| GFP (Av-GFP JL-8) | Living colors | 1:10,000 |
| Total RB1 | Cell signaling | 1:1,000 |
| BMI1 | Millipore | 1:5,000 |

**Table S8. Plasmid list used in oncogenic senescence induction studies.**

| Plasmid | Description |
|---|---|
| **pMam-neo** | Control vector containing dexamethasone inducible MMTV promoter driving expression of Neomycin resistance gene |
| **pMam-mp15** | Mouse p15 expression vector with MMTV inducible promoter and Neo resistance |
| **pMam-EGFP-*ink4ab*** | Zebrafish *ink4ab* fused to EGFP and inserted into pMam vector |
| **pEGFPc1** | Control plasmid containing CMV promoter and EGFP |
| **pEGFP-ink4ab** | Zebrafish *ink4ab* cloned into pEGFPc1 |
| **pCR3-mp15** | Mouse mp15 expressed from CMV promoter |
| **pBabe-puro** | Control plasmid with puromycin resistance |
| **pBabeHRas$^{V12}$** | Oncogenic HRas$^{V12}$ cloned into pBabe |
